# The Intestinal Microbiome, Dietary Habits, and Physical and Psychological Resilience in Postpartum Women

**DOI:** 10.1101/2022.09.07.506896

**Authors:** Michiko Matsunaga, Mariko Takeuchi, Satoshi Watanabe, Aya K. Takeda, Takefumi Kikusui, Kazutaka Mogi, Miho Nagasawa, Keisuke Hagihara, Masako Myowa

**Affiliations:** Department of Advanced Hybrid Medicine, Graduate School of Medicine, Osaka University, Osaka, Japan; Graduate School of Education, Kyoto University, Kyoto, Japan; Japan Society for the Promotion of Science, Tokyo, Japan; Cykinso, Inc., Tokyo, Japan; School of Veterinary Medicine, Azabu University, Kanagawa, Japan

## Abstract

The population of postpartum women suffering from mental illness is increasing steadily, particularly under conditions of the COVID-19 pandemic. Identifying factors that contribute to resilience in postpartum women is urgently needed to decrease risks of poor physical and psychological functioning. Studies have linked variations in the intestinal microbiota to depression in clinical samples, but the impacts in postpartum women in a Japanese population are unknown. We conducted two studies to examine the links between intestinal microbiota, physical condition, and psychological state in nonclinical, postpartum Japanese women. Our results show that decreasing *Lachnospira* and alpha diversity of microbiome is related to high mental health risk (i.e., parenting stress and/or depression). Psychological resilience and physical conditions were associated with relative abundances of genera *Blautia, Clostridium, Eggerthella*. This study contributes to further understanding of the gut-brain axis mechanisms and supports proposals for interventions to enhance resilience in postpartum women.

## Introduction

In developed countries, 10%–15% of postpartum women suffer from depression ^1, 2^, and proportions have increased as a result of the COVID-19 pandemic; the latest data in Japan show that 28.7% of postpartum women have high risk of depression ^3^. Postpartum depression can persist for long periods ^4-6^, and prolonged depression can affect children’s mental health and cognitive development ^7, 8^. Preventing mental illnesses such as depression and parenting stress among postpartum women requires identifying not only risk factors but also factors that enhance resilience.

Recent studies have shown that the intestinal microbiota is associated not only with physical diseases (e.g., diabetes and cancer) but also with psychological disorders (e.g., depression and anxiety) ^9, 10^. For example, a recent systematic review found that patients with depression and anxiety disorders have increased inflammatory gut bacteria and relatively few short-chain fatty acid-producing gut bacteria ^11^. The gut microbiota affects brain functions through autonomic nerves, neuropeptides, hormones, and the immune system ^12^. Understanding the relationships between the intestinal microbiota and mental health in postpartum women will allow for developing mental health prevention and intervention methods that utilize the gut microbiota (e.g., administering probiotics or improving dietary habits) ^13^.

In this study, we focus on Japanese postpartum women. It is important to note that the composition of the intestinal microbiota in the Japanese population differs from that in Western and even other Asian countries (e.g., China) ^14^. In particular, *Blautia* is a dominant bacterium in the intestinal microbiota of Japanese people, and researchers have found that it is related to functional components of traditional Japanese fermented foods ^15^. We considered it necessary to examine how the intestinal microbiota relate to Japanese postpartum women’s physical and mental health, considering the Japanese food culture and dietary habits.

For Study 1, we analyzed 347 stool samples of Japanese postpartum women caring for infants and toddlers from 0 to 4 years old, and investigated factors-- including intestinal microbiota-- associated with mental illness (i.e., depression and parenting stress). In Study 2, we specifically focused on 27 primiparous women in the early postpartum period (within three to six months). We examined the relationships among their intestinal microbiota, physical and physiological functions, and psychological resilience.

## Results

### Study 1: Relationships between mental health risk and physical condition, dietary and lifestyle habits, and intestinal microbiota among postpartum women raising children aged 0∼4 years

We assessed mental health risk using the Parenting Stress Index (PSI) and the Beck Depression Inventory (BDI-II), and defined the high-risk group as women whose scores exceeded the cutoff values determined in the manual (see *Questionnaires* in Supplementary Information for more details on cutoff settings). Although no participants in this study had been diagnosed with any physical or psychological diseases, 82 of the 347 postpartum women (23.63%) were at high risk for mental illness (i.e., parenting stress and/or depression) (Figure 1a). Although high parenting stress increases the risk of depression, notably, not all mothers above the parenting stress cutoff were above the depression cutoff. As we cannot rule out the possibility that intestinal microbiota, physical condition, and lifestyle factors associated with stress and depression risks may differ, we compared stress risk and depression risk separately with a healthy control group (Figure 1b).

**Figure 1.**
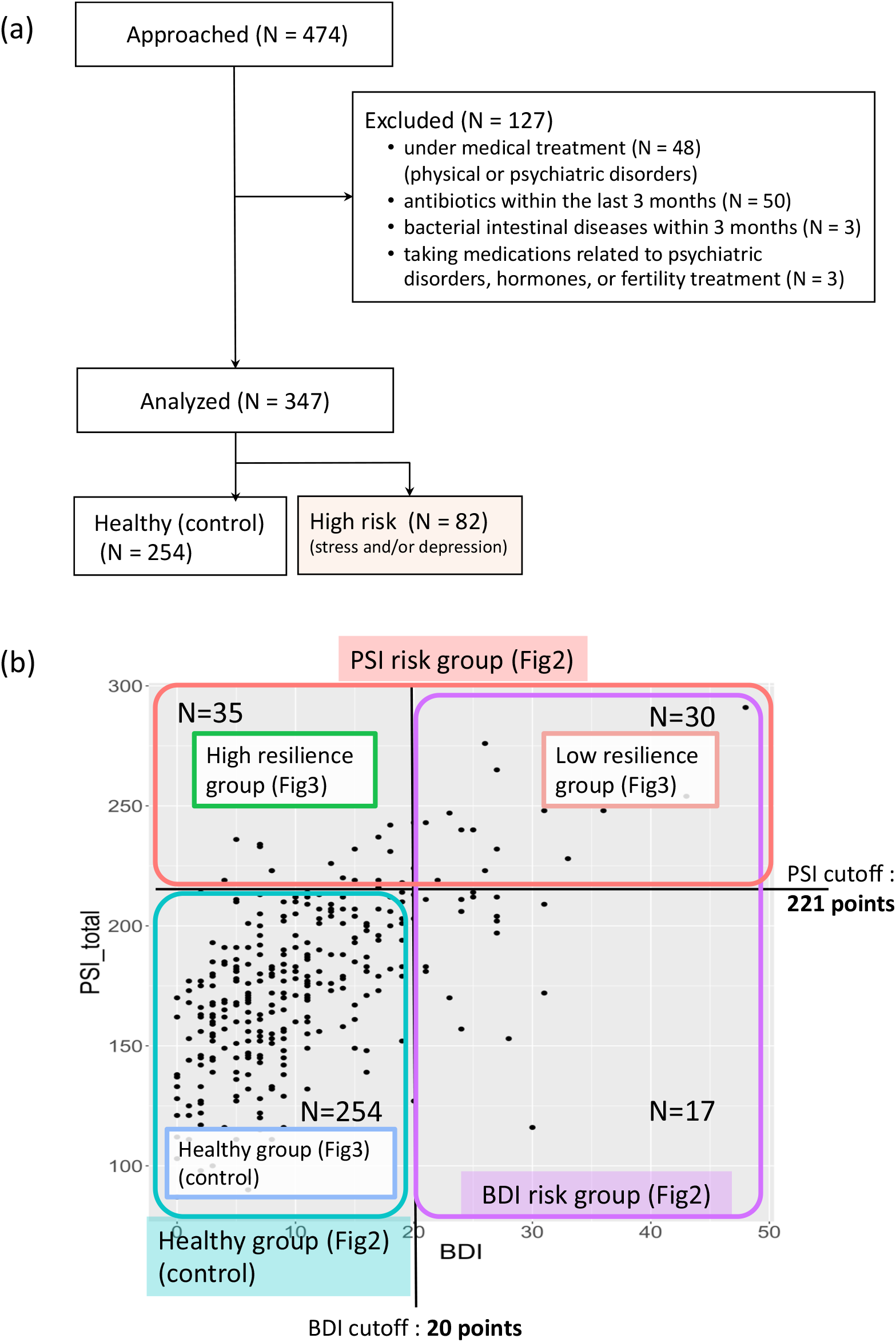
Study 1 consort diagram and classification of psychological risk in mothers. (a) A consort flow diagram of Study 1. We finally analyzed data from 347 postpartum Japanese women data. We evaluated the participants’ mental health using the parenting stress questionnaire (PSI) and the depression questionnaire (BDI) with their standardized cutoff scores. A total of 254 participants were classified as the healthy (control) group and 82 as the risk group (i.e., risk of stress and/or depression). (b) The scatterplot shows an example of distribution of parenting stress (PSI) and depression (BDI) scores. The standardized cutoff for PSI is a total score of 221 points, with sub-scales 124 points for the parental aspect and 101 points for the child aspect. Participants exceeding any of the PSI cutoff values were classified in the parenting stress risk group. The standardized cutoff for the BDI was 20 points, and participants exceeding the BDI cutoff were classified in the depression risk group. Participants below the cutoff for both PSI and BDI were classified as the healthy group (see *Questionnaires* in Supplementary Information for details). The analysis shown in Figure 2 compares the healthy group to the parenting stress risk group and to the depression risk group. The analysis shown in Figure 3 compares three groups: healthy group, high resilience group (i.e., high parenting stress but no depression), and low resilience group (i.e., high parenting stress and depression).

Regarding physical condition, dietary and lifestyle habits, we found that women at high risk for both parenting stress and depression had lower subjective sleep quality ratings and worse physical condition scores than those of the healthy group (*p* <.01, *q* <.01) (Table 1). Especially for physical condition, all of the Multi-dimensional Physical Scale (MDPS) scores were related to both high parenting stress and depression risk, that is, digestive system dysfunction, physical depression, and low female hormone function-impaired microcirculation was related to risk of parenting stress and depression. In terms of diet, the frequencies of milk and cheese consumption were lower in both risk groups (*p* <.05, *q* <.05) (Table 1). In addition, the depression risk group showed lower staple food intake than did the healthy group (*p* =.03, *q* =.09).

**Table 1.**
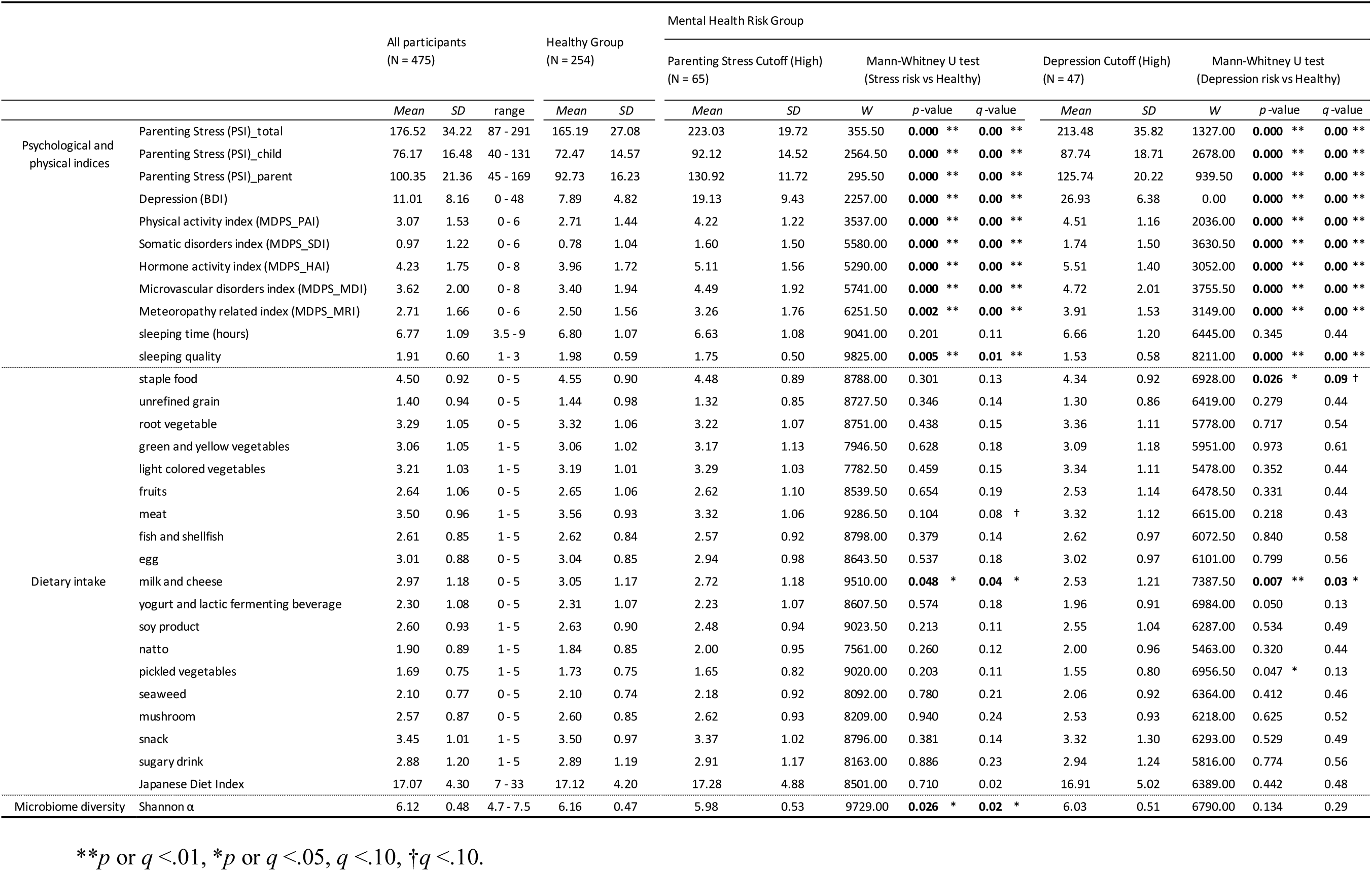
Comparison between the Mental Health Risk and Healthy Groups (Study 1)

In analyzing the intestinal microbiota we found less *Lachnospira* in the depression risk group than in the healthy group (*p* =.01, *q* =.04) (Figure 2, Supplementary Table 5). Regarding the parenting stress risk, we found lower alpha diversity in the stress risk group than in the healthy group (*p* =.03, *q* =.02), and there were significant group differences in the following bacteria: *Odoribacter, Alistipes, Lachnospira, Monoglobus, Phascolarctobacterium, Veillonella, Sutterella, Escherichia-Shigella*. (Figure 2, Supplementary Table 5).

**Figure 2.**
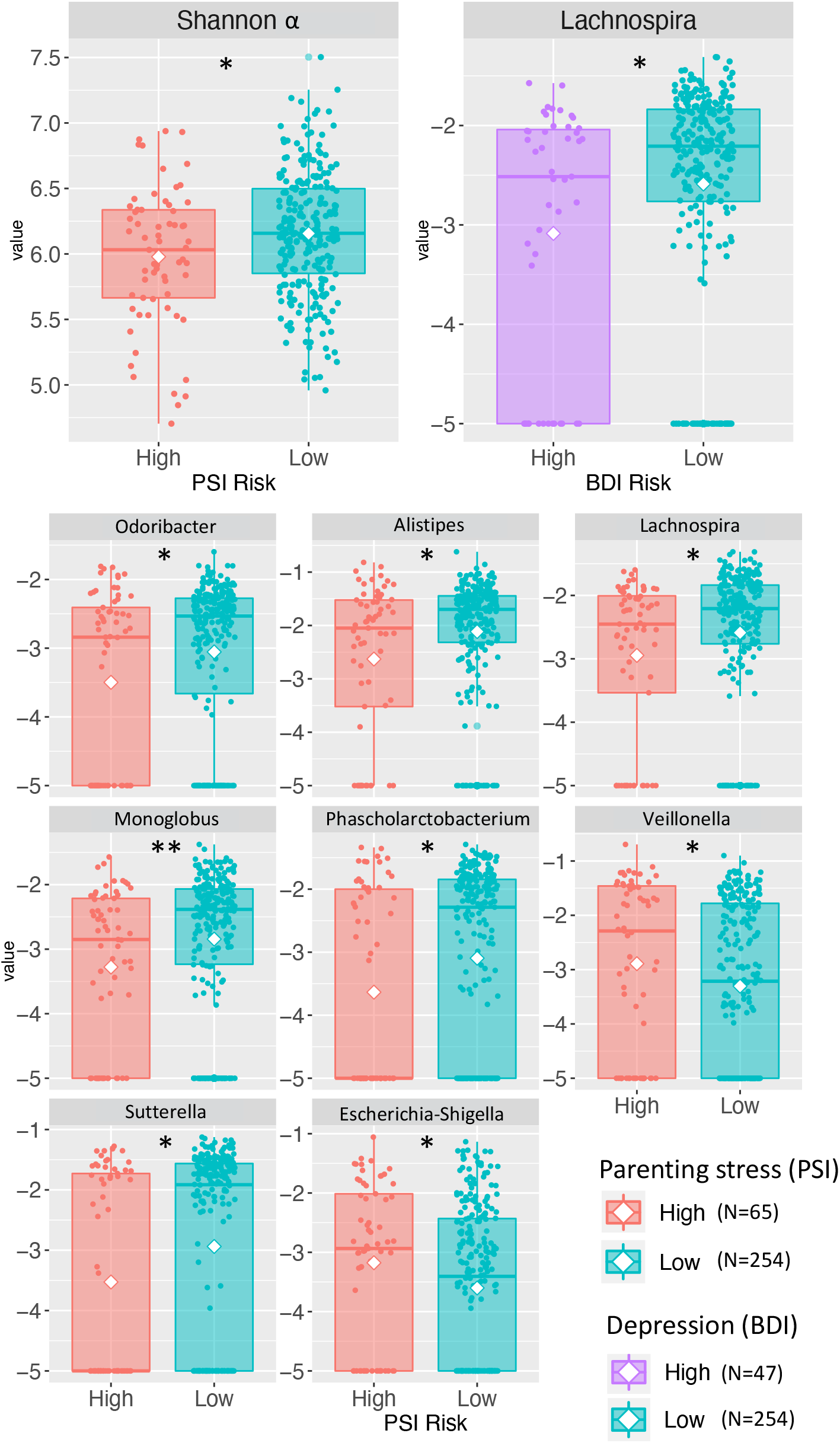
Parenting stress and depression score comparisons between healthy and high-risk groups (Study 1). *\*\*p* <.01, *\*p* <.05.

### Post-hoc analysis in Study 1

#### Psychological resilience and the Japanese diet

The women in the parenting stress risk group were classified into one of the two categories: those whose PSI and BDI scores exceeded the cutoff (n =30, 8.65% of the total) and those whose PSI scores exceeded the cutoff but their BDI scores did not (n = 35, 10.09% of the total) (Figure 1b). That is, we observed the same high child-rearing stress in both groups, but we considered that the women who were not severely depressed were more resilient: The former group had low resilience, whereas the latter group had high resilience. To clarify the factors that contributed to psychological resilience, we compared the healthy group with the high and low-resilience groups.

We identified significant group differences in multigroup comparisons for both total PSI score and parent aspect of PSI score, BDI score, almost all physical condition scores (MDPS), and sleep quality score (*p* <.01, *q* <.01), but no significant difference in child aspect of PSI score (Figure 3). Notably, the low-resilience group showed significantly lower indices of physical activity (PAI), hormone activity (HAI), microvascular disorders (MDI), and meteropathy (MRI) than the other two groups. This finding suggests that physical condition is a potential factor associated with psychological resilience. Regarding the intestinal microbiota, there were significant group differences in *Odoribacter, Lachnoclostridium, Flavonifractor, Monoglobus*, and *Phascolarctobacterium* (*p* <.05, *q* <.06). However, only *Flavonifractor* showed a significant group difference between the high- and low-resilience groups (Figure 3). Supplementary Table 6 presents the detailed statistics.

**Figure 3.**
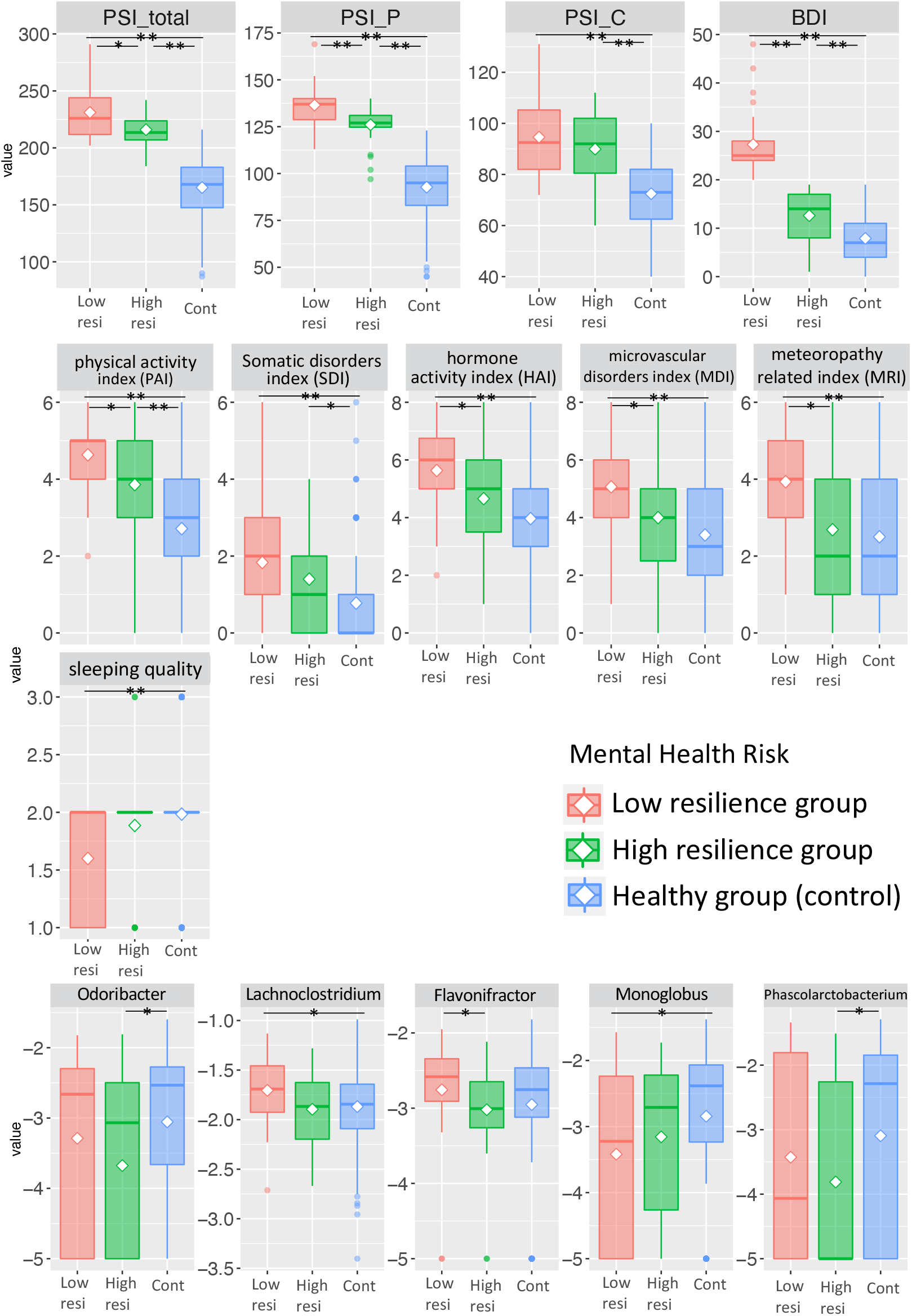
Steel-Dwass test results for comparisons of resilience factors (physical conditions, dietary and lifestyle habits) with intestinal microbiota (Study1). Low resi: low-resilience group (both PSI and BDI above the cutoff); High resi: high resilience group (PSI above cutoff, BDI below cutoff); Cont: healthy group (both PSI and BDI below cutoff). *\*\*p* <.01, *\*p* <.05.

To examine the effects of the Japanese diet on postpartum women’s mental health and physical condition, we administered the Japanese Diet Index (JDI) and compared high- and low-score groups using the median as the cutoff. These analyses revealed that the high JDI group had lower scores on BDI (*p* =.04, *q* =.09), PAI (related to digestive system function; *p* =.03, *q* =.06), and MDI (i.e., less impaired microcirculation related to female hormonal function; *p* =.02, *q* =.06). Regarding intestinal microbiota, the high JDI group had more *Agathobacter* and *Subdoligranulum* than the low JDI group (*p* =.03, *q* =.06; *p* =.02, *q* =.06, respectively) (Figure 4). (See Supplementary Table 7 for detailed statistics.)

**Figure 4.**
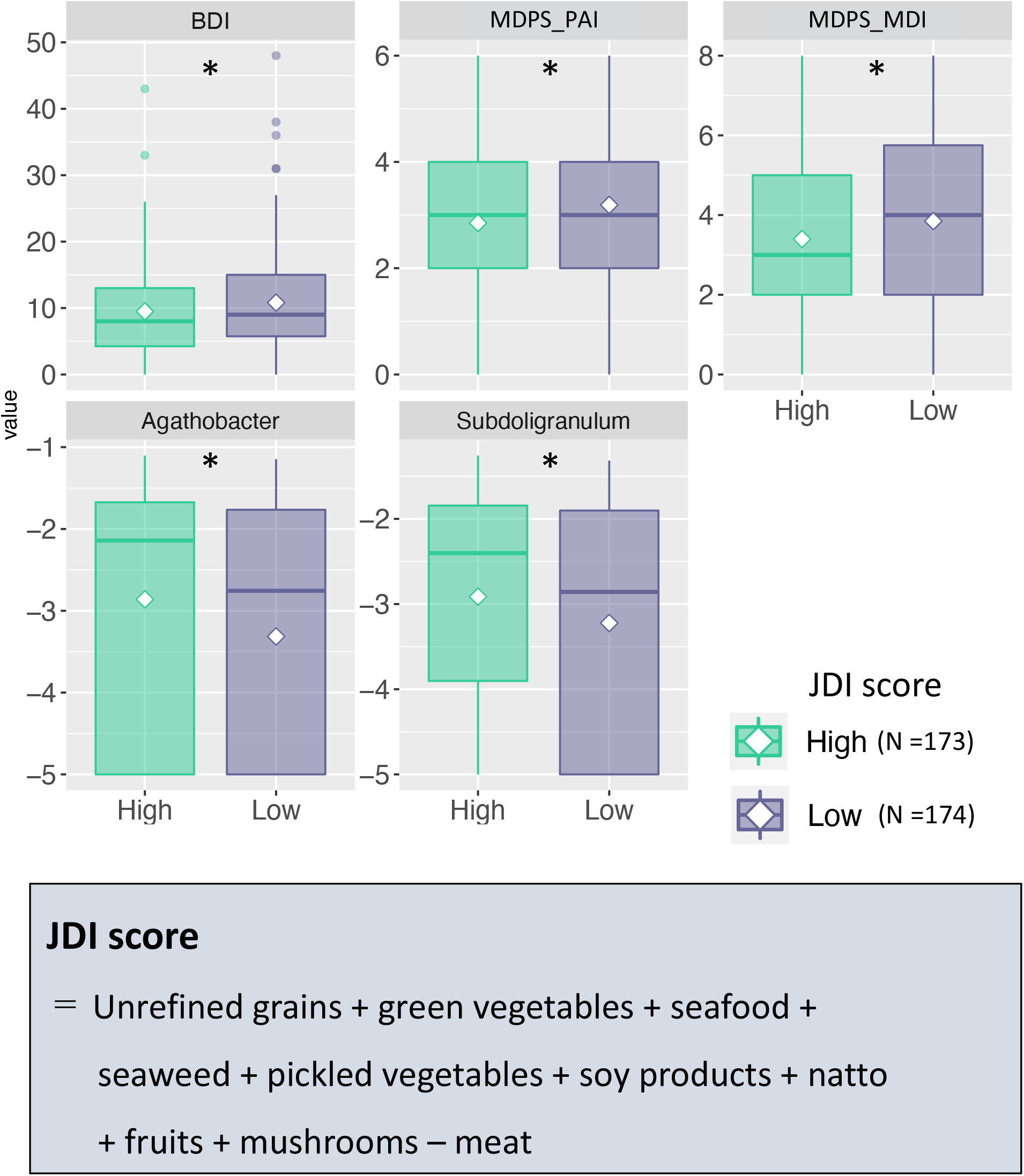
Comparison between high and low Japanese dietary intake scores groups (Study 1). PAI: physical activity index, MDI: microvascular disorders index *\*p* <.05

### Study 2: Body composition and physical function of women in the early postpartum period

To more clearly understand Japanese women’s physical condition in the early postpartum period, we referred to medical diagnostic criteria or reference values for women of the same age from previous studies (Table 2). Figure 5a shows that body mass index (BMI) was similar to the reference values for most of our participants. However, skeletal muscle mass index (SMI) was lower than the medically diagnosed criterion for sarcopenia in nearly half of the participants (n = 13, 40.74%; Figure 5b). Figure 5c shows lower hand grip strength than the reference values for most participants (n = 23, 85.19%); hand grip strength is considered to reflect total body muscle strength. Furthermore, Figures 5d, 5e, and 4f show lower values for most participants than reference values for the two-step test for overall lower limb motor function (n = 26, 96.30%), normal gait speed (n = 19, 70.37%), and maximum gait speed (n = 25, 92.59%). These results indicate lower muscle mass and poorer motor function among Japanese postpartum women than among control women of the same age.

**Table 2.** Body Composition and Physical Function among Japanese Postpartum Women and Reference Values (Study 2)

|  | *Mean* | *SD* | Min | Max | Reference value |  |
| --- | --- | --- | --- | --- | --- | --- |
| mother's age (years) | 33.63 | 4.18 | 27 | 41 | - |  |
| mother's education (years) | 16.19 | 1.39 | 12 | 19 | - |  |
| BMI (kg/m<sup>2</sup>) | 20.54 | 2.67 | 15.4 | 26.80 | 21.6 kg/m<sup>2</sup> | Ministry of Health, Labour and Welfare, 2016<sup>16</sup> |
| FAT (%) | 26.49 | 5.68 | 14.9 | 37.30 | 24.70% | data provided by Inbody, Inc. |
| SMI (kg/m<sup>2</sup>) | 5.89 | 0.59 | 5.00 | 7.20 | < 5.7 kg/m<sup>2</sup> | diagnostic criteria of sarcopenia |
| ECW/TBW | 0.383 | 0.004 | 0.373 | 0.390 | 0.382 | Ohashi et al., 2018<sup>17</sup> |
| handgrip (kg) | 24.05 | 4.21 | 16.10 | 32.80 | 28.77 kg | Japan Sports Agency, 2017<sup>18</sup> |
| two-step test | 1.37 | 0.14 | 0.82 | 1.54 | 1.53 | Yamada et al., 2020<sup>19</sup> |
| normal gait speed (m/s) | 1.18 | 0.20 | 0.70 | 1.64 | 1.26 m/s | Obuchi et al., 2020<sup>20</sup> |
| maximum gait speed (m/s) | 1.92 | 0.38 | 1.08 | 3.00 | 2.5-2.8 m/s | Yokohama Sports Medicine Center, 2015<sup>21</sup> |
| RS25 | 89.44 | 8.46 | 74 | 107 | - |  |
| J-RS | 93.93 | 13.61 | 69 | 115 | - |  |
| PSI_parent | 172.67 | 35.76 | 111 | 231 | - |  |
| PSI_infant | 98.89 | 22.32 | 53 | 144 | - |  |
| PSI_total | 73.78 | 19.03 | 46 | 108 | - |  |
| CESD | 7.81 | 4.59 | 0 | 18 | - |  |
| CVI | 4.25 | 0.25 | 3.77 | 4.62 | - |  |
| CSI | 2.95 | 0.82 | 1.4 | 4.33 | - |  |
| oxytocin (pg/ml) | 129.41 | 93.86 | 25.57 | 409.82 | - |  |

Reference values other than for SMI are averages for women of the same age.

**Figure 5.**
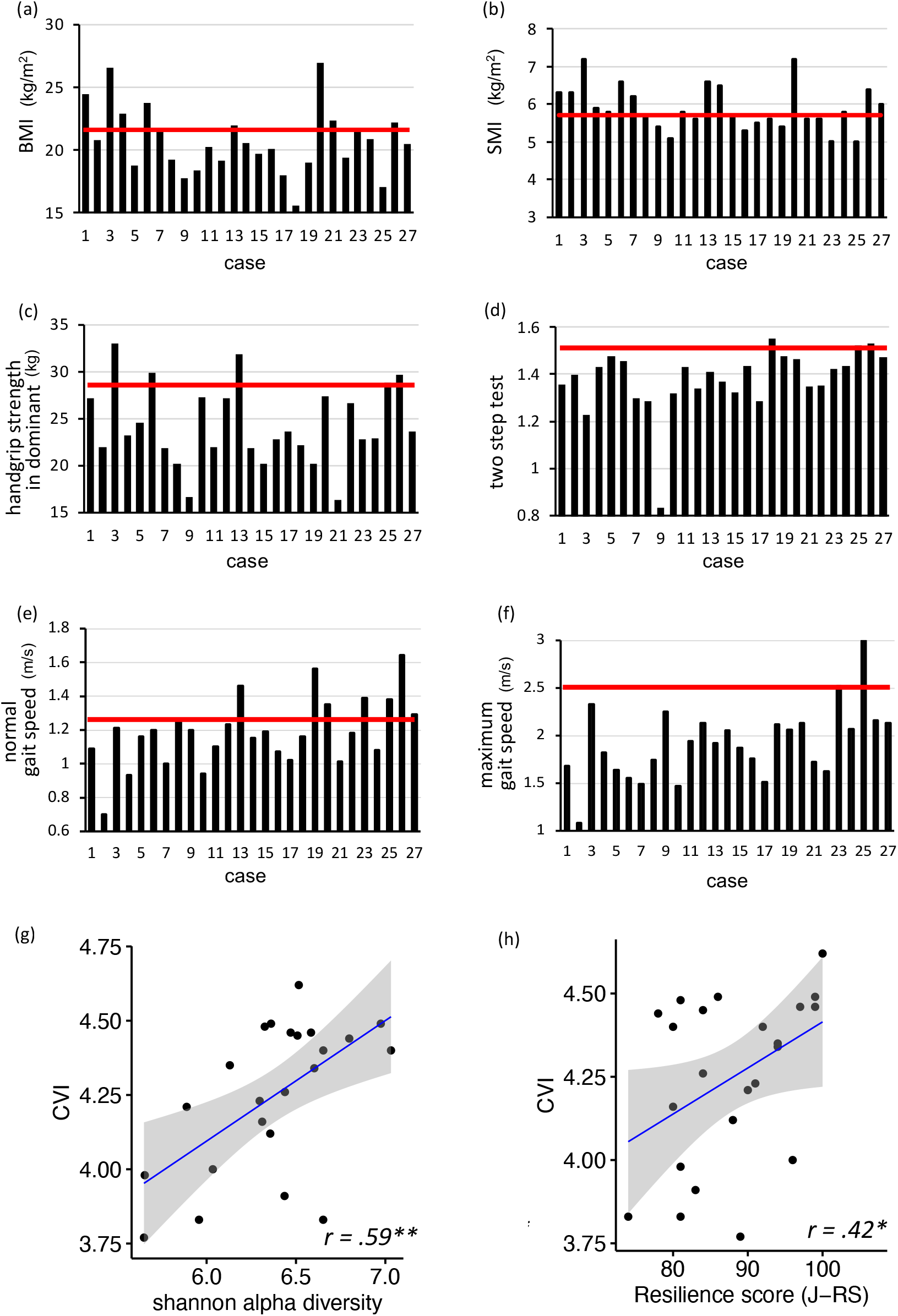
Physical assessment of women in the early postpartum period (Study2). (a)-(f) show physical condition of each participant. The horizontal axis (i.e., case) and black bars indicate representative values for each participant. Red lines indicate the reference values shown in Table 2. (g) Pearson’s correlation between vagal nervous function (CVI) assessed by electrocardiogram and intestinal microbiota (Shannon α diversity). (f) Pearson’s correlation between CVI and resilience (J-RS) scores assessed by questionnaires. Gray bands in (g) and (f) show 95% confidence intervals, and each dot represents a single participant. \**\*p* <.01, *\*p* <.05.

### Intestinal microbiota, autonomic nervous system functioning, and resilience in the early postpartum period (Study 2)

We investigated the relationships between intestinal microbiota, physical and physiological function, and psychological resilience among Japanese women in the early postpartum period. First, we found a positive correlation between Shannon α and vagal nervous activity (*r* =.59, *p* =.003, 95% confidence interval [CI; 0.005–0.71]) (Figure 5g and Supplementary Fig. 2). In addition, vagal nervous activity was positively correlated with psychological resilience (J-RS) (*r* =.42, *p* =.047, 95% CI [0.23–0.80]) (Figures 5h and Supplementary Fig. 1).

Next, shotgun metagenome analysis and regression analysis of the women’s core microbiota showed that *Blautia SC05B48, Clostridium SY8519, Collinsella aerofaciens*, and *Eggerthella lenta* significantly explained the individual differences in psychological resilience and physical function (Figure 6, Supplementary Figs. 2 and 3, Supplementary Table 8). Specifically, *Blautia SC05B48* was positively associated with psychological resilience (J-RS) and two-step test score (Figures 6a and b). *Clostridium SY8519* was positively associated with psychological resilience (J-RS) and negatively associated with oxytocin (Figures 6c and d). *Colinsella aerofaciens* was positively associated with psychological resilience (J-RS) and negatively associated with maximum gait speed (Figures 6e and f). *E. lenta was* negatively associated with psychological resilience (RS25), and positively associated with parenting stress (child aspect subscale) and dominant hand grip strength (Figures 6g, h, and i). Finally, *Faecalibacterium prausnitz* was positively associated with high sympathetic nerve activity among autonomic functions (Figure 6j). Supplementary Fig. 3 shows the results for identifying bacteria by shotgun metagenomic analysis, and Supplementary Table 8 shows the null results including statistics.

**Figure 6.**
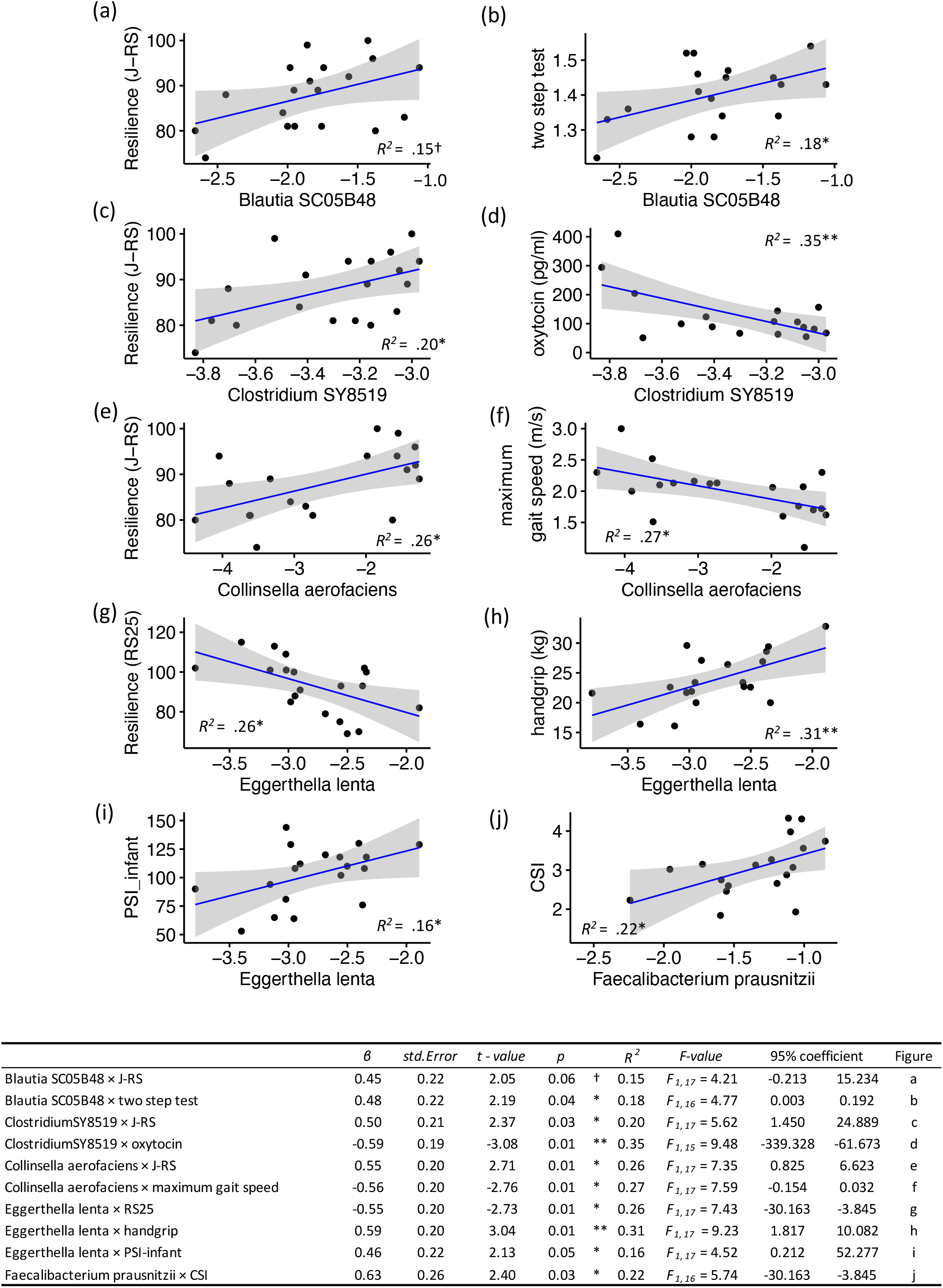
Relationships between individual differences in intestinal microbiota and physical and psychological resilience. Scatter plots and regression lines with 95% confidence intervals show significant relationships between intestinal microbiota (i.e., horizontal axis) and physical and psychological indices (i.e., vertical axis). Each dot represents a single participant. In the regression analysis of *Blautia SC05B48* and the two-step test, we excluded one participant from the statistical analysis because her two-step test score was an outlier (mean minus 3 *SD* or less); PSI_infant = child aspect of parenting stress index (subscale of PSI), CSI = the cardiac sympathetic nervous index, *β* = standardized partial regression coefficient, ** *p* <.01, * *p* <.05, † *p* =.06 (effect size > medium (*R*^*2*^ >.13)).

## Discussion

A particularly noteworthy new finding from these two studies is that of associations between three new factors and psychological resilience in Japanese postpartum women: the intestinal microbiota (e.g., Shannon-αdiversity, *Lachnospira, Blautia, Eggerthella*), physical conditions related to female hormone function, and autonomic nervous system function (e.g., vagal nervous activity).

In Study 1 we examined factors associated with mental health risk in a large sample of postpartum Japanese women caring for young children aged 0–4 years. We found that approximately 24% of the women were at high risk for mental illness, based on their PSI and BDI scores, although no participants were diagnosed with clinical psychiatric or physical illness. Postpartum women with high parenting stress and depression were commonly in poorer physical condition including hormonal imbalance and poor digestive function and blood circulation. In particular, postpartum women with high depression scores reported notably reduced sleep quality and less frequent consumption of staple foods than did the postpartum women in the healthy group; this result may be in accordance with the DSM-V diagnostic criteria for depression, insomnia and anorexia. *Lachnospira* is a butyrate-producing bacterium that has many beneficial effects on the intestinal environment, such as improving the immune function of the intestinal mucosa and inhibiting cancer cells ^22^. The most recent systematic review also found fewer butyrate-producing bacteria in depressed patients ^11^.

The high parenting stress risk group had a less diverse intestinal microbiota (Shannon α) than healthy postpartum women, with three notable features in their core bacteria: fewer gut bacteria related to immunity and antibiotic effects (e.g., *Odoribacter, Sutterella, Lachnospira*), more inflammatory gut microbiota (e.g., *Escherichia-Shigella*), and abnormal levels of acetic acid- and propionic acid-producing bacteria (e.g., *Veillonella, Alistipes, Phascolarctobacterium*). Regarding the first and second features,previous studies found that *Odoribacter* produced antibacterial “isoalloLCA,” which suppresses the growth of intestinal bacteria that cause diarrhea and abdominal pain ^23^.

Studies in mice have shown that *Sutterella* was also associated with alpha-defensins, a type of immunity in the gut that fluctuates in response to psychological stress ^24^. In contrast, *Escherichia-Shigella* is a recognized major pathogen that causes diarrhea, and it is associated with the severity of anxiety disorders ^25^. These findings suggest that the postpartum Japanese women with high parenting stress would also show poorer intestinal epithelial barrier function and increased intestinal inflammation.

Regarding the third gut feature above, the hydrogen produced in fermenting dietary fiber is mainly consumed to produce acetic acid in the Japanese gut, whereas in Europe, the United States, and China, it is consumed for methane production ^15^. Acetic acid suppresses inflammation in the intestine and promotes repair of epithelial cell damage, and propionic acid contributes to energy and lipid metabolism in the colon ^26, 27^. However, some researchers reported that excessive acetic acid and propionic acid increased psychosomatic diseases and stress: higher concentrations of acetic acid and propionic acid produced by *Veillonella* were related to severity of irritable bowel syndrome ^28^. In summary, in our results, abnormal intestinal microbiota and metabolites appeared to cause physical problems and nerve inflammation, and postpartum women who are at high risk for parenting stress may experience excessive psychological stress.

In Study 2 we focused on primiparous postpartum women within six months of giving birth and examined factors contributing to their physical and psychological resilience. We found that vagal nerve activity was positively related to both diversity of intestinal microbiota (Shannon α) and psychological resilience (J-RS). High vagal function at rest reflects capacity for stress tolerance ^29^, and the vagal nerve is less active under stressful situations and associated with stress hormones such as cortisol ^30^. To our knowledge, there has been only one examination of a relationship between vagal activity and resilience, a study of trauma in US military personnel ^31^. We also showed that individual differences in vagal activity were a possible biomarker of psychological resilience; the diversity of the gut microbiota could contribute to psychological resilience through individual differences in vagal activity.

Among the core intestinal microbiota, we found that *Blautia SC05B48, Clostridium SY8519, Collinsella aerofaciens*, and *Eggathella lenta* explained individual differences in both psychological resilience and physical function in primiparous women in the early postpartum period. *Blautia* and *Clostridium* are butyrate-producing bacteria, and *Blautia* in particular is more prevalent in Japanese than in Western and other Asian populations ^14^. It has attracted attention for its potential positive effects on physical health (e.g., metabolic syndrome, lifestyle-related diseases, and healthy life expectancy) ^32^. The results of the present study provide new evidence for relationships between individual differences in *Blautia* and psychological resilience in postpartum Japanese women.

In addition, *Clostridium* contributes to suppressing intestinal inflammation and allergic responses by inducing the production of regulatory Treg cells that suppress immune system responses ^33^. In the present study, *Clostridium SY8519* was positively related to J-RS score but negatively to oxytocin. We explain this finding as a function of oxytocin and the stress system. Under stressful conditions, the hypothalamic-pituitary-adrenal (HPA) axis is activated to secrete cortisol, and at the same time, the oxytocin system in the hypothalamic paraventricular nucleus is activated^34, 35^. In another study, oxytocin secretion was shown to suppress cortisol secretion ^36^. One interpretation of our results is that postpartum women with high *Clostridium SY8519* and low oxytocin have little physiological need for immune responses or stress responses on the HPA axis to restore homeostasis. It is widely accepted that higher oxytocin levels in new mothers have positive effects on the mother and child ^37-39^. Future studies should aim to examine and clarify the relationships between intra-individual variations in oxytocin, stress responses, and intestinal microbiota.

*E. lenta* was negatively correlated with psychological resilience (RS25) in this study, and positively correlated with parental stress and grip strength, consistent with previous findings of increased *Eggerthella* in patients with major depressive disorder ^40^. An important feature of *E. lenta* is its ability to metabolize isoflavones to produce equol, which has female hormone-like effects ^41, 42^. Equol has many benefits for women’s health, including protection against breast cancer, osteoporosis, and arteriosclerosis. The Japanese diet features high soybean intake, but only about 50% of the population has equol-producing microbes in the gut; in the United States and European countries, the proportions are around 20%–30% ^42^.

*Clostridium SY8519* is known to be involved in the process of metabolizing isoflavones ^43^, and inthe post-hoc analysis performed for Study 1, we also found that both female hormone function and increased *Flavonifractor* were associated with resilience. *Flavonifractor* is known to be involved in the degradation of flavonoids, which have antioxidant, stress-reducing, anticancer, and immunological properties. Indeed, in previous studies, excess *Flavonifractor* decreased the bioavailability of flavonoids and correlated negatively with depression and physical quality of life ^10, 44^. Isoflavones are among the flavonoids, and the Japanese diet traditionally includes a variety of foods that are rich in flavonoids (e.g., green tea, soybeans, sesame, *yuzu* (a small citrus fruit)). Therefore, the Japanese diet and its associated gut microbiota metabolism could contribute to physical recovery among postpartum women.

Consistent with our finding that Japanese food may contribute toward the physical and psychological health of postpartum women, experiments on mice have revealed that glucosylceramide, a component of *koji*, increased *Blautia* ^15^; *koji* is an ingredient used in most Japanese traditional fermented foods (e.g., soy sauce, miso, sake, and vinegar). Several researchers have reported that dietary intervention with glucosylceramide derived from soybean and rice bran reduced colon, head, and neck cancers ^45, 46^.With further research, Japanese food has the potential to serve a prebiotic function of improving intestinal microbiota in postpartum recovery.

Regarding physical functioning, in Study 2 we found that approximately 41% of the primiparous postpartum women had SMI of less than 5.7 kg/m^2^, which met one of the criteria for sarcopenia. Upper extremity muscle strength and lower extremity motor function were also lower in most of the mothers than the reference values for women of the same age. SMI peaks at around age 20–30 and then declines by 1%– 2% per year ^47^, and age-related decline in SMI can ultimately lead to frailty and sarcopenia. Therefore, it is important to assess the degree of decline and recovery of muscle mass and physical functions from pre-pregnancy to postpartum.

There were several limitations in our study. In particular, we collected the data for Study 2 before the wider impacts of COVID-19 had hit. Although we had planned a large sample size, we had to discontinue the study and conduct preliminary analyses based on data from only 27 cases. Furthermore, we conducted multiple comparisons to examine the relationships among intestinal microbiota, physical and physiological condition, and psychological state, but because of the small sample size we were unable to perform *p*-value correction or mediation analysis. While we acknowledge the possibility of multiple correlations, all of our results had medium to large effect sizes, and the single regression analysis also confirmed an association. We have also reported all statistics with nonsignificant results and scatter plots in detail. Another limitation is that because both studies were cross-sectional, we could not evaluate intrapersonal changes before and after childbirth. Longitudinal studies with larger samples are also required to clarify neurophysiological-psychological relationships by including samples from clinical groups with high levels of stress and depression.

In summary, we investigated relationships among the following four variables; intestinal microbiota, autonomic nervous system function, physical condition (e.g., body composition, physical function and physical condition) and psychological state. Figure 7 showed a putative conceptual model of physical and psychological resilience. The results of the present study show that intestinal microbiota and autonomic nervous system function are related to physical and psychological resilience in postpartum women. Indeed, we discovered that physical condition as assessed by the MDPS questionnaire (e.g., weak digestive system function and female hormone function) is a key factor in whether psychological resilience is maintained and remains the same as in the pre-vulnerable state (Figure 7b) or whether it falls into the vulnerable state (Figure 7c). Poorer physical conditions are related to low psychological resilience, that is, a high risk of both depression and parenting stress. The intestinal microbiota of postpartum women in this vulnerable state included decreased Shannon-αdiversity and butyrate-producing bacteria (e.g.,*Lachnospira*), suggesting that the intestinal environment is in an inflammatory state (Figure 7c). Weakened autonomic nervous system function and physical function could impede recovery and lead to prolonged physical and mental illness. It is important to screen and intervene with postpartum women who are at risk in the pre-vulnerable state (Figure 7b), to enhance their resilience before they become vulnerable (Figure 7c). In terms of intervention, the Japanese diet and the predominant gut microbiota in the Japanese population (e.g., *Blautia*) were associated with psychological and physical resilience in this study. As it is widely known that the composition of intestinal microbiota differs among ethnic populations ^14, 15^, the association of intestinal microbiota and resilience have some ethnic-specific aspects. This suggests that effective support and health care for depression and stress should be developed in a tailor-made manner taking into account ethnic traits and individual differences. In conclusion, our study contributes to both further understanding of the microbiota-gut-brain axis ^12^ underlying individual differences in psychological disorders, and proposing personalized interventions that can enhance resilience.

**Figure 7.**
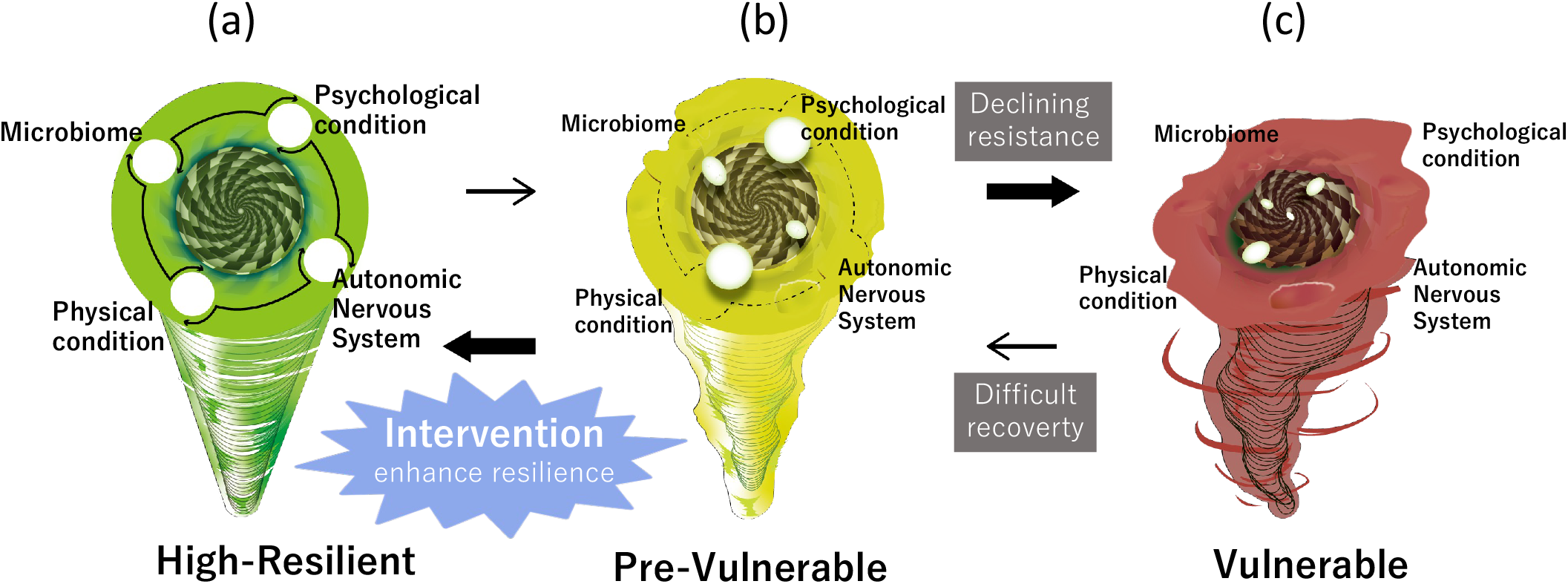
Putative conceptual model of physical and psychological resilience. (a) High resilient state in which all physical and psychological factors are healthy. (b) Pre-vulnerable state, characterized by the onset of decline in physical and psychological functions, before falling into the vulnerable state. (c) Vulnerable state, with deterioration of physical and psychological conditions.

## Methods

### Participants and Procedures

#### Study 1

Study 1 is part of a Japanese research project entitled *The Principle of Human Social Brain-Mind Development*. For this portion of the project, we analyzed data from 347 participants (mean age = 34.62 years, *SD* = 4.81, range: 21 to 47 years) (Figure 1a). No participants had any psychological or physical disorders. We collected the data between January and February 2021. For each participant, we obtained written informed consent and collected a stool sample, and then gave them four questionnaires to complete at home: PSI, BDI-II, Multidimensional Physical Scale (MDPS, physical condition), and Mykinso Pro(dietary and lifestyle habits). We also collected information on socioeconomic status. This study was approved by the Medical Ethics Committee of Kyoto University (no. R2624) and registered in the UMIN system (UMIN000042508). See *Participants’ demographic information* and *Questionnaires* in Supplementary Information for more detailed information on participants characteristics (Supplementary Table 1) and the questionnaires.

#### Study 2

Study 2 is part of the Japanese research project entitled *Investigation of the Interrelationship among Mental, Physical and Intestinal Microbiota in Postpartum Mothers and Their Infants*. For this study, we analyzed data from 27 first-time mothers within three to six months after childbirth (mean age = 33.63 years, *SD* = 4.18, range 27 to 41 years). None of the participants had any diagnosed psychiatric disorders. We collected the data between September 2019 and March 2020. All participants visited The Baby Laboratory at Kyoto University for the study and provided written informed consent. This study was approved by the Medical Ethics Committee of Osaka University (no. 18409 and no. 20074) and the Ethics Committee of Kyoto University (no. 27-P-1) and registered in the UMIN system (UMIN000045125).

Participants visited the lab twice. At the first visit, we collected the physical data and administered the questionnaires: the Japanese version of the Resilience Scale (RS25 and J-RS, psychological resilience) and the Center for Epidemiological Studies-Depression scale (CESD). The physical measurements used to assess physical composition and function were as follows: (1) BMI, body fat percentage, SMI, and extracellular water/total body water ratio (ECW/TBW) (InBody 770 multi-frequency bioelectrical impedance device, InBody Japan Inc., Tokyo); (2) grip strength of the dominant hand, which is considered to reflect the muscular strength of the whole body; (3) the two-step test for risk of locomotive syndrome; a lower score indicates weak gait function including muscle strength, balance, and flexibility of the lower limbs; (4) normal and maximum gait speed for 5 m; lower gait speed score also indicates declining motor function in the lower limbs.

At the second visit, we collected saliva samples to measure oxytocin, collected three minutes of resting ECG data to assess the autonomic nervous system (i.e., cardiac sympathetic and cardiac vagal index; based on Toichi et al., 1997 ^48^), administered the PSI, and collected information on age, years of education, and breastfeeding schedule. Although we asked the participants to visit twice within the same week if they could, the mean interval between the two visits was 7.74 days (*SD* = 8.19 days). To evaluate the intestinal microbiota, participants collected stool samples at home within two or three days after the second visit. See *Participants’ demographic information, Questionnaires, Physical and physiological measurements in Study2* in Supplementary Material for more details.

### Gut-microbiome analysis (Study 1 and Study 2)

#### Fecal sample collection and DNA extraction

Fecal samples were collected using Mykinso fecal collection kits containing guanidine thiocyanate solution (Cykinso, Tokyo, Japan), transported at ambient temperature, and stored at 4°C. DNA extraction from the fecal samples was performed using an automated DNA extraction machine (GENE PREP STAR PI-480, Kurabo Industries Ltd, Osaka, Japan) according to the manufacturer’s protocol.

#### 16S rRNA gene sequencing

The detailed sequencing methods are described elsewhere ^49^. Briefly, amplicons of the V1V2 region were prepared using the forward primer (16S_27Fmod: TCG TCG GCA GCG TCA GAT GTG TAT AAG AGA CAG AGR GTT TGA TYM TGG CTC AG) and the reverse primer (16S_338R: GTC TCG TGG GCT CGG AGA TGT GTA TAA GAG ACA GTG CTG CCT CCC GTA GGA GT). The libraries were sequenced in a 250-bp paired-end run using the MiSeq Reagent Kit v2 (Illumina; 500 cycles).

#### Metagenomic shotgun sequencing (Study 2)

DNA for each sample was sheared to about 300 bp using the Covaris ME220 (Covaris, Woburn, MA, USA). The shotgun sequencing library was constructed using the KAPA Hyper Prep Kit (Roche). Libraries were sequenced on the DNBSEQ-G400 (BGI, Shenzhen, China) at 150 bp paired end.

#### Bioinformatics Analysis

##### (1) 16S rRNA analysis (Study 1)

The data processing and assignment using QIIME2 pipeline (version 2020.8) ^50^ was performed as described elsewhere ^51^.

##### (2) 16S rRNA analysis (Study 2)

The data processing and assignment based on the QIIME2 pipeline (version 2019.4) ^50^ were performed in the following steps: (1) DADA2 ^52^ for joining paired-end reads, filtering, and denoising, and (2) assigning taxonomic information to each ASV using a naive Bayes classifier in the QIIME2 classifier with the 16S rRNA gene V1V2 region data from Greengenes ^53^ to determine identity and bacterial composition.

##### (3) Metagenomic shotgun analysis (Study 2)

Each read pair was mapped against a custom database including human genome (hg19), bacteria and fungi in RefSeq, and viruses in nucleotide collection (NCBI-nt) by Kraken2 ^54^.

##### (4) Statistical Analysis

We performed all data manipulation, analyses, and graphics using R and RStudio (versions 3.5.1 and 1.1.456, respectively) and used the R package qiime2R and microbiome R for all analyses. We used the R package tidyMicro (version 1.48) and ggplot2 for visualizations. We measured alpha diversity at the ASV level using the Shannon index. In addition, we analyzed the microbiota shared by the mothers (so-called phylogenetic common core ^55^) using the categorized postpartum age group data. We determined the composition of the core microbiota to be the set of genus-level bacterial groups shared by at least 50% of the participants in relative abundance (RA) of at least 0.01%; we calculated RA as the number of sequencing reads of each taxon in a sample standardized by the total number of sequences generated for that sample. We only included in the analyses taxa that were present in at least one sample, and we aggregated sequence counts for taxa that did not meet these requirements into the “other” category. We applied these filters at the phylum, family, and genus levels, and left sequence counts that we could not classify to the taxonomic level of interest as unclassified counts of the lowest level possible. See Supplementary Tables 3 and 4 for more details of the core microbiome detected in the Studies 1 and 2.

### Statistical Analysis

We performed statistical analyses and visualizations for both studies using R and RStudio (versions 4.0.5 and 1.4.1106, respectively). We performed *q*-value estimation with R and RStudio (versions 4.2.1 and 1.4.1106, respectively) using R package Bioconductor (version 3.15). All statistical tests were used two-tailed test.

#### Study 1

We defined the high-risk group as women whose PSI and/or BDI scores exceeded the cutoff scores for parenting stress and/or depressive symptoms; we defined healthy women as those whose scores were below the cutoffs. We used the Mann-Whitney *U* test to compare the high-risk and healthy groups for physical state score, dietary and lifestyle habits, and intestinal microbiota (i.e., alpha diversity and the core microbiota). We calculated false discovery rate-adjusted *q* for all indices. We performed two post-hoc statistical analyses for Study 1; for both, we set significance at *p* <.05 and *q* <.10.

##### (1) Resilience factors

We observed two categories of the parenting stress risk group, women whose PSI and BDI scores both exceeded the cutoffs (n =30, 8.65% of the total) and women whose PSI score exceeded the cutoff but whose BDI score did not (n = 35, 10.09% of the total). We observed the same high child-rearing stress in both groups, but considered the women who were not severely depressed to be more resilient than the severely depressed women. Therefore, for the post-hoc analysis, we focused on the high parenting stress risk group and divided them into high (i.e., high parenting stress and low depression, n = 35) or low (i.e., high parenting stress and high depression, n =30) resilience. To clarify factors contributing to resilience, we compared women in the high parenting stress risk group with the healthy group (control). First, we used the Kruskal-Wallis test to compare the three groups, then for all variables that were significant at p <.05 we made multiple comparisons with the Steel-Dwass test. We set significance at *p* <.05 and *q* <.10.

##### (2) Effects of the Japanese diet

Previous researchers reported evidence that the Japanese diet was associated with risk of physical diseases and dementia ^56, 57^. However, intervention studies also showed that consuming a Japanese diet for one month could change intestinal microbiota and improve health ^55^. To examine the effects of Japanese food intake on mental and physical health and intestinal microbiota, we also calculated the JDI score and divided the subjects at the median into high and low JDI groups; we then compared the groups with Welch’s t-test, with significance set at *p* <.05 and *q* <.10. See *Japanese Dietary Intake (Study 1)* in Supplementary Information for more detailed information on calculating the JDI score.

#### Study 2

In this study we aimed to investigate relationships among intestinal microbiota, physical and physiological function, and psychological resilience in early postpartum women. First, we conducted an exploratory correlation analysis and focused on the variables for which *p* <.05 or with a moderate effect size (*r* ≥.30). Next, to clarify the extent to which individual differences in intestinal microbiota could explain individual differences in psychosomatic states, we constructed a regression model. The objective variable was the RA of gut bacteria identified by the shotgun metagenomic analysis, and the dependent variables were the psychological, physical, and physiological index scores. As this was an exploratory study with a small sample size, we set significance of the regression model at p <.05 or when the effect size of the model was above moderate (R^2^ ≥.13).

## Acknowledgements

We thank Daisuke Motooka (Genome Information Research Center, Research Institute for Microbial Diseases, Osaka University) for conducting the shotgun metagenomic analysis. This study was supported by a Grant-in-Aid for Scientific Research from Japan Society for the Promotion of Science (JSPS)(17H01016 to M. Myo. and 19K21813 to M. Myo.); a Grant-in-Aid for JSPS Fellows (19J15173 and 22J01448 to M. M.); and a grant from the Center of Innovation Program, Japan Science and Technology Agency (JPMJCE1307 to M. Myo.).

## Authors’ contributions

Conceptualization: M.M., M.T., K.H., M.Myo.; statistical analysis: M.M., T.M., S.W.; behavior assessment and analysis: M.M., T.M.; microbiome analysis: S.W., A.T.; hormone analysis: T.K., K.M., M.N.; writing−original draft: M.M., T.M.; writing−review & editing: M.M., T.M., S.W., A.T., T.K., K.M.,M.N., K.H., M.Myo.; visualization: M.M., M.T.; supervision: M.Myo., K.H.; funding acquisition: M.Myo., K.H., T.K.

## Data Availabiliry

All sequencing data has been deposition to the NCBI Sequence Read Archive under project and are publicly available as of the date of the publication. The BioProject accession numbers are follow: no. PRJNA844514 (16 s rRNA gene sequencing) for Study1, no. PRJNA844516 (16 s rRNA gene sequencing) and no. PRJNA###### (Metagenomic shotgun sequencing, now preparing) for Study 2. All of the questionnaire data and physiological data (i.e., physical composition, autonomic nervous system, oxytocin hormone) are available from the Figshare database (DOI: 10.6084/m9.figshare.20439837).

