## Supplementary Information for "The Intestinal Microbiome, Dietary Habits, and Physical and Psychological Resilience in Postpartum Women"

#### Participant demographic information (Study 1 and Study 2)

##### *Study 1*

Study 1 is part of a research project in Japan entitled *The Principle of Human Social Brain-Mind Development*. This project aims to create a database of intestinal microbiota of more than 1,000 mother-child pairs in Japan and to investigate the relationships among those microbiota and emotional and cognitive development. Participants were recruited through nursery schools throughout Japan that agreed to cooperate in the study. Study 1 reports only data on intestinal microbiota collected between January and February 2021.

Figure 1a in the main text showed the consort diagram of the data collected on 474 postpartum women who were raising children aged 0–4 years in nursery schools. Of these 474 participants we excluded 127 for the following reasons: (i) having a physical or psychological disease under medical treatment (e.g., diabetes, irritable bowel syndrome, depression) at the time of data collection, (ii) having taken antibiotics within the last three months, (iii) having had a bacterial intestinal disease within the last three months, and (iv) taking medications related to psychiatric disorders, hormones, or fertility treatments (i.e., Lexapro, Sertraline, Lutoral, Lithium carbonate, levothyroxine sodium hydrate). Finally, we analyzed data from 347 participants (mean age = 34.62 years, *SD* = 4.81, range: 21 to 47 years).

Supplementary Table 1 shows the characteristics of Study 1 participants. All had completed at least nine years of education, and annual household income ranged from less than 3000,000 yen (about 26,000 USD) to more than 15,000,000 yen (about 130,000 USD). Around 52% of respondents reported annual income of less than 7,000,000 yen (about 60,700 USD). The average household income of Japan's child-rearing generation in 2019 was 7,459,000 yen (about 64,700 USD) [1]. That is, more than 60% of the participants in Study 1 had below-average annual household incomes.

**Supplementary Table 1. Characteristics of Study 1 Participants**

| | *Mean* | *SD* | range |
|---|---|---|---|
| Mother's age (years) | 34.62 | 4.81 | 21 - 47 |
| Education（years） | 14.61 | 1.99 | 9 - 22 |
| **Houeshold income** | | **N** | **%** |
| less than 3 million yen (about 26,000 USD) | | 38 | 10.95 |
| 3-5 million yen (about 26,000 - 43,360 USD) | | 88 | 25.36 |
| 5-7 million yen (about 43,360 - 60,700 USD) | | 93 | 26.80 |
| more than 7 million yen (about 60,700 USD) | | 104 | 29.97 |
| no answer | | 24 | 6.92 |
| **Number of children** | | **N** | **%** |
| one | | 89 | 21.33 |
| two or more | | 244 | 67.72 |
| Pregnant with second child | | 24 | 6.92 |
| no answer | | 14 | 4.03 |
| **Age of youngest child (Years since last childbirth)** | | **N** | **%** |
| less than 1 year old | | 43 | 12.39 |
| 1-2 years old | | 158 | 45.53 |
| more than 3 years old | | 129 | 37.18 |
| no answer | | 17 | 4.90 |

***Study 2***

Study 2 is part of the Japanese research project entitled *Investigation of the Interrelationship among Mental and Physical Health and Intestinal Microbiota in Postpartum Mothers and Their Infants*. We collected and analyzed data from 27 first-time postpartum women within three to six months after childbirth (mean age = 33.63 years, *SD* = 4.18, range: 27–41 years). The average postpartum duration was 152.52 days (*SD* = 19.10 days, range: 115–191 days). None of the participants had any diagnosed psychiatric disorders. All 27 participants continued breastfeeding their infants during the study period: 15 women (55.56%) breastfed exclusively, and 12 (44.44%) continued breastfeeding supplemented with formula. Twenty-two women (81.48%) gave birth vaginally, and five (18.52%) gave birth by cesarean section. Eleven women (40.74%) used antimicrobials during delivery. We collected data between January and February 2021.

### Questionnaires

#### *Study 1*

We administered four questionnaires during Study 1 to measure the new Japanese mothers' parenting stress, depression, physical condition, and dietary habits. We measured parenting stress using the Japanese version of the Parenting Stress Index (PSI), a questionnaire developed in the United States [2] and validated and standardized in Japanese [3]. The PSI consists of 78 items, and respondents rate each item on a 5-point scale. It two subscales: stress related to the child's characteristics (i.e., the child aspect) and stress related to the parents themselves (i.e., the parent aspect). The cutoff scores to indicate high parenting stress are 221 points for total score, 101 points for child aspects, and 124 points for parent aspect.

We measured depression using the Beck Depression Questionnaire (BDI-II) [4], which assesses the severity of depressive symptoms. The BDI is based on the diagnostic criteria of the DSM-IV and has been validated and standardized in Japanese [5]. It consists of 21 items rated on 4-point scales. Based on the treatment guidelines of the Japanese Society of Mood Disorders, we considered 20 points as the cutoff for depression because the urgency of medical intervention and treatment increases with scores higher than 20.

To assess physical condition, we used the Multidimensional Physical Scale (MDPS) (Takeuchi et al., 2022, under review), which is based on Oriental medicine criteria for physical pathology in postpartum women. The MDPS consists of 17 items on five subscales, with each item rated on a 3-point scale (Supplementary Table 2); the five subscales are physical activity index (PAI) (higher scores indicate physical inactivity [e.g., tiredness, physical lethargy]); (ii) somatic disorders index (SDI) (higher scores indicate high physical depression symptoms [e.g., anorexia, bloating]); (iii) hormone activity index (HAI) (higher scores indicate poor (female) hormone function [e.g., coldness, dizziness, dry skin]), (iv) microvascular disorders index (MDI) (higher scores indicate impaired microcirculation related to (female) hormone function [e.g., skin pigmentation and rough skin]); and (v) meteoropathy-related index (MRI) (higher scores indicate impaired water metabolism and meteorological diseases [e.g., swelling, headache due to bad weather]).

We measured dietary and lifestyle habits using the Mykinso Pro Questionnaire, which includes questions on lifestyle (smoking, drinking, sleep, defecation), dietary intake, and physical diseases (history of medical treatment, medication). The questionnaire was developed by Cykinso Co., a provider of intestinal microbiota screening services in Japan. For sleep time and sleep quality, participants reported their average number of hours of sleep per day, and sleep quality on a 3 point scale (3 points: good, 2 points: fine, 1 point: bad). For dietary intake, the questionnaire comprises 18 components: staple foods (e.g., rice, bread and noodles), unrefined grain, root vegetables, green and yellow vegetables, light colored vegetables, fruits, meat, fish and shellfish, egg, milk and cheese, yogurt and lactic acid bacteria beverage, soybeans and soybean-derived foods, natto (fermented soybeans), seaweed, pickles, mushroom, snack, and sugary drinks. The questionnaire asks participants how frequently they consumed each food during the last week of stool collection, using the following 5-point scale as follows: 0 points: did not eat, 1 point: 1–3 times/week, 2 points: 4–6 times/week, 3 points: once/day, 4 points: twice or more/day). The questionnaire also collected demographic information (age, years of education, etc.) and socioeconomic status.

**Supplementary Table 2. Multidimensional physical scale (MDPS)**

The following is a list of the ways you might have felt. Please select the number for how often you have felt this way recently

| | | | 0: none of the time or rarely | 1: occasionally or sometimes | 2: most or all of the time |
|---|---|---|---|---|---|
| PAI | MDPS 1 | I get tired more easily than before. | ☐ | ☐ | ☐ |
| | MDPS 2 | I am sleepy after a meal. | ☐ | ☐ | ☐ |
| | MDPS 3 | I feel sluggish in my hands or legs. | ☐ | ☐ | ☐ |
| SDI | MDPS 4 | I have a sensation of a lump or foreign body in the throat. | ☐ | ☐ | ☐ |
| | MDPS 5 | I feel that I haven't completely emptied my bladder after urination. | ☐ | ☐ | ☐ |
| | MDPS 6 | I feel bloated. | ☐ | ☐ | ☐ |
| HAI | MDPS 7 | My hands and legs feel cold. | ☐ | ☐ | ☐ |
| | MDPS 8 | My hair loss has become serious. | ☐ | ☐ | ☐ |
| | MDPS 9 | I suffer from dry skin. | ☐ | ☐ | ☐ |
| | MDPS 10 | I suffer from vaginal itchiness or dryness. | ☐ | ☐ | ☐ |
| MDI | MDPS 11 | I have bags under my eyes. | ☐ | ☐ | ☐ |
| | MDPS 12 | I have age spots. | ☐ | ☐ | ☐ |
| | MDPS 13 | I have a rough skin. | ☐ | ☐ | ☐ |
| | MDPS 14 | I have hemorrhoids. | ☐ | ☐ | ☐ |
| MRI | MDPS 15 | I have headaches. | ☐ | ☐ | ☐ |
| | MDPS 16 | I feel dizzy. | ☐ | ☐ | ☐ |
| | MDPS 17 | I have swelling. | ☐ | ☐ | ☐ |

Abbreviations: MDPS, Multidimensional Physical Scale; PAI, physical activity index; SDI, somatic disorders index; HAI, hormone activity index; MDI, microvascular disorders index; MRI, meteoropathy-related index.

***Study 2***

We administered five questionnaires in Study 2 to assess parenting stress, depression, and psychological resilience, and demographic information. We measured parenting stress with the PSI as in Study 1. We measured depression in Study 2 using the Center for Epidemiological Studies-Depression scale (CESD), which was developed by the National Institute of Mental Health [6] and standardized for Japanese [7]. The CESD consists of 20 items rated on 4-point scales, and the cutoff for depression is 16 points.

To assess psychological resilience, we used the Resilience Scale (RS25) [8, 9] and a Japanese version of the scale (J-RS) developed by Takeuchi et al. (2022, under review). The RS25 is a validated, reliable, and popular quantitative resilience assessment scale that is widely used in Europe and the United States. It consists of 25 items rated on 7-point scales. Researchers have attempted to validate a Japanese version of the RS25, but it did not have the same reliability or validity as the original [9]. The researchers concluded that the scale might not be suitable for Japanese people because the latter tend to refrain from positive and proactive expressions. Other resilience scales developed in Japan, such as the Mental Resilience Scale and the Two-Dimensional Resilience Factors Scale, have only been validated on students or young adults. For this study, in addition to the RS25, we used the J-RS, which includes items that take into account the

Japanese characteristic of placing importance on empathy and social connections; it has been validated in adults (Takeuchi et al., 2022, under review).

**Japanese Dietary Intake (Study 1)**

Researchers have identified nine components of a traditional Japanese dietary pattern: rice, miso, seafood, green and yellow vegetables, seaweed, pickles, green tea, beef/pork, and coffee [10]. Recently proposed is the Modern Japanese Diet Score (JDI12), which adds another three beneficial components: soybeans and soy-derived foods, fruits, and mushrooms [11]. We did not include coffee or green tea in the JDI score because the Mykinso Pro questionnaire did not measure these. The questionnaire asked respondents how frequently they had eaten each food component during the week of stool collection using the following 5-point scale: 0 point: did not eat; 1 point: 1–3 times/week; 2 points: 4–6 times/week; 3 points: once/day; 4 points: twice or more/day. We then calculated the dietary intake score by applying respondents' scores to the 4-point scales in the following formula JDI12 ＝ Unrefined grains + green vegetables + seafood + seaweed + pickled vegetables + soy products + natto +  mushrooms + fruits + mushrooms – meat.

**Physical and physiological measurements in Study 2**

***Body composition and physical function***

We conducted four types of tests to assess body composition and physical function among Japanese postpartum women who were new mothers: body composition itself, grip strength, the two-step test, and gait speed. Body composition was assessed using the InBody 770 multi-frequency bioelectrical impedance (BIA) device (InBody Japan Inc., Tokyo, JPN). Participants stood on a balance scale, grasped the handle of the machine, and remained still for 1 minute while maintaining appropriate posture. We collected the following data from the BIA device: body weight, BMI, body fat percentage, skeletal muscle mass index (SMI), and extracellular water/total body water ratio (ECW/TBW). We calculated BMI, body fat percentage, and SMI according to the following formulas:

- *Body Mass Index (BMI) (kg/m$^2$) = weight (kg)/square of height (m$^2$)*
- *body fat percentage = fat mass (kg)/body weight (kg) × 100*
- *Skeletal Muscle Mass Index (SMI (kg/m$^2$) = skeletal muscle mass (kg)/square of height (m$^2$).*

We measured grip strength in the dominant hand using a digital grip strength meter (TKK-5401, Matsuyoshi Medical Instrument Co., Ltd., Tokyo, JPN). The participants grasped a steering wheel device with their arms stretched out to the side and lowered. We measured both hands twice, alternating between them, and used the highest value for the dominant hand for the analysis.

In the two-step test, we measured the distance between two steps taken with as large a stride as possible and calculated the score using the following formula:

*2Step test score = maximum stride length (cm)/height of individual (cm).*

We took two measurements and used the average for each individual.

We measured maximum and normal gait speed for each participant, using a stopwatch to measure the time required to walk a distance of 5 m on an 11-m walking course with 6 m for acceleration and deceleration on a flat floor. We measured maximum gait speed as the speed at which the participant was walking as quickly as she could, and normal gait speed as when she walked as her normal pace. We took each measurement twice, and used the average speed for normal gait speed, and the faster time of the two for maximum gait speed.

#### *Autonomic nervous system function*

To measure individual differences in autonomic nervous system function, we recorded electrocardiograms (ECG) using the Polymate Mini AP108 (Miyuki Giken Co., Ltd., Tokyo, JPN) at 1000 Hz with electrodes attached at the V2 induction of the three-point induction. The R-R interval was calculated on the last minute of ECG data during the 3-min resting period with infants. We analyzed the cardiac vagal and sympathetic indices following a method that applied Lorenz plot analysis [12]. Lorenz plots are more robust than frequency analysis for analyzing ECG data in a short period of time or in the presence of noise due to body movement. In addition, they have the advantage of allowing for calculating sympathetic and vagal activities separately and independent of respiration and posture.

#### *Hormone data collection and analysis*

Prior to saliva collection, participants rinsed their mouths with water. After 10 minutes, they chewed on an oral cotton swab (Salimetrics, State College, PA, USA) placed sublingually for 3 minutes to collect saliva. We immediately stored the collected saliva at −80 °C, and repeated the process to obtain a backup sample. We used the first samples to calculate each individual's hormone levels. However, if the first sample contained insufficient saliva or if it had outlier hormone levels, we used the second sample.

We measured oxytocin concentration with a commercial kit (ADI-900-153A-0001; Enzo Co. Ltd., Tokyo, JPN) following the manufacturer's protocol. In brief, we dried 240 μL of saliva using a Speedvac evaporator at room temperature for 3 h and reconstituted it in 240 μL of assay buffer; we then used 100 μL of that for the assay.

### Supplementary Table 3. The Core Microbiome in Study 1

| core_genus | Kingdom | Phylum | Class | Order | Family | Genus |
|---|---|---|---|---|---|---|
| Actinomyces | Bacteria | Actinobacteria | Actinobacteria | Actinomycetales | Actinomycetaceae | Actinomyces |
| Agathobacter | Bacteria | Firmicutes | Clostridia | Clostridiales | Lachnospiraceae | Agathobacter |
| Alistipes | Bacteria | Bacteroidetes | Bacteroidia | Bacteroidales | Rikenellaceae | Alistipes |
| Anaerostipes | Bacteria | Firmicutes | Clostridia | Clostridiales | Lachnospiraceae | Anaerostipes |
| Bacteroides | Bacteria | Bacteroidetes | Bacteroidia | Bacteroidales | Bacteroidaceae | Bacteroides |
| Bifidobacterium | Bacteria | Actinobacteria | Actinobacteria | Bifidobacteriales | Bifidobacteriaceae | Bifidobacterium |
| Bilophila | Bacteria | Proteobacteria | Deltaproteobacteria | Desulfovibrionales | Desulfovibrionaceae | Bilophila |
| Blautia | Bacteria | Firmicutes | Clostridia | Clostridiales | Lachnospiraceae | Blautia |
| Butyricicoccus | Bacteria | Firmicutes | Clostridia | Clostridiales | Ruminococcaceae | Butyricicoccus |
| Colidextribacter | Bacteria | Firmicutes | Clostridia | Clostridiales | Oscillospiraceae | Colidextribacter |
| Collinsella | Bacteria | Actinobacteria | Coriobacteriia | Coriobacteriales | Coriobacteriaceae | Collinsella |
| Dorea | Bacteria | Firmicutes | Clostridia | Clostridiales | Lachnospiraceae | Dorea |
| Eggerthella | Bacteria | Actinobacteria | Coriobacteriia | Coriobacteriales | Coriobacteriaceae | Eggerthella |
| Erysipelatoclostridium | Bacteria | Firmicutes | Bacilli | Erysipelotrichales | Erysipelatoclostridiaceae | Erysipelatoclostridium |
| Escherichia-Shigella | Bacteria | Proteobacteria | Gammaproteobacteria | Enterobacteriales | Enterobacteriaceae | Escherichia-Shigella |
| Faecalibacterium | Bacteria | Firmicutes | Clostridia | Clostridiales | Ruminococcaceae | Faecalibacterium |
| Flavonifractor | Bacteria | Firmicutes | Clostridia | Clostridiales | Oscillospiraceae | Flavonifractor |
| Fusicatenibacter | Bacteria | Firmicutes | Clostridia | Clostridiales | Lachnospiraceae | Fusicatenibacter |
| Intestinibacter | Bacteria | Firmicutes | Clostridia | Clostridiales | Clostridiaceae | Intestinibacter |
| Lachnoclostridium | Bacteria | Firmicutes | Clostridia | Clostridiales | Lachnospiraceae | Lachnoclostridium |
| Lachnospira | Bacteria | Firmicutes | Clostridia | Clostridiales | Lachnospiraceae | Lachnospira |
| Lachnospiraceae_UCG-004 | Bacteria | Firmicutes | Clostridia | Clostridiales | Lachnospiraceae | UCG-004 |
| Monoglobus | Bacteria | Firmicutes | Clostridia | Monoglobales | Monoglobaceae | Monoglobus |
| Negativibacillus | Bacteria | Firmicutes | Clostridia | Clostridiales | Ruminococcaceae | Negativibacillus |
| Odoribacter | Bacteria | Bacteroidetes | Bacteroidia | Bacteroidales | Marinifilaceae | Odoribacter |
| Oscillibacter | Bacteria | Firmicutes | Clostridia | Clostridiales | Oscillospiraceae | Oscillibacter |
| Parabacteroides | Bacteria | Bacteroidetes | Bacteroidia | Bacteroidales | Tannerellaceae | Parabacteroides |
| Parasutterella | Bacteria | Proteobacteria | Gammaproteobacteria | Burkholderiales | Sutterellaceae | Parasutterella |
| Phascolarctobacterium | Bacteria | Firmicutes | Negativicutes | Acidaminococcales | Acidaminococcaceae | Phascolarctobacterium |
| Romboutsia | Bacteria | Firmicutes | Clostridia | Clostridiales | Clostridiaceae | Romboutsia |
| Roseburia | Bacteria | Firmicutes | Clostridia | Clostridiales | Lachnospiraceae | Roseburia |
| Sellimonas | Bacteria | Firmicutes | Clostridia | Clostridiales | Lachnospiraceae | Sellimonas |
| Streptococcus | Bacteria | Firmicutes | Bacilli | Lactobacillales | Streptococcaceae | Streptococcus |
| Subdoligranulum | Bacteria | Firmicutes | Clostridia | Clostridiales | Ruminococcaceae | Subdoligranulum |
| Sutterella | Bacteria | Proteobacteria | Gammaproteobacteria | Burkholderiales | Sutterellaceae | Sutterella |
| Turicibacter | Bacteria | Firmicutes | Bacilli | Erysipelotrichales | Erysipelotrichaceae | Turicibacter |
| Tuzzerella | Bacteria | Firmicutes | Clostridia | Clostridiales | Lachnospiraceae | Tuzzerella |
| UBA1819 | Bacteria | Firmicutes | Clostridia | Clostridiales | Ruminococcaceae | UBA1819 |
| Veillonella | Bacteria | Firmicutes | Negativicutes | Veillonellales | Veillonellaceae | Veillonella |

### Supplementary Table 4.  The Core Microbiome in Study 2

| core_genus | Kingdom | Phylum | Class | Order | Family | Genus |
|---|---|---|---|---|---|---|
| Actinomyces | Bacteria | Actinobacteria | Actinobacteria | Actinomycetales | Actinomycetaceae | Actinomyces |
| [Eubacterium] | Bacteria | Firmicutes | Erysipelotrichi | Erysipelotrichales | Erysipelotrichaceae | [Eubacterium] |
| Adlercreutzia | Bacteria | Actinobacteria | Coriobacteriia | Coriobacteriales | Coriobacteriaceae | Adlercreutzia |
| Clostridiaceae_Clostridium | Bacteria | Firmicutes | Clostridia | Clostridiales | Clostridiaceae | Clostridium |
| Lachnospiraceae_Clostridium | Bacteria | Firmicutes | Clostridia | Clostridiales | Lachnospiraceae | Clostridium |
| Ruminococcus | Bacteria | Firmicutes | Clostridia | Clostridiales | Ruminococcaceae | Ruminococcus |
| Erysipelotrichaceae_Clostridium | Bacteria | Firmicutes | Erysipelotrichi | Erysipelotrichales | Erysipelotrichaceae | Clostridium |
| Bacteroides | Bacteria | Bacteroidetes | Bacteroidia | Bacteroidales | Bacteroidaceae | Bacteroides |
| Barnesiella | Bacteria | Bacteroidetes | Bacteroidia | Bacteroidales | [Barnesiellaceae] | Barnesiella |
| Bifidobacterium | Bacteria | Actinobacteria | Actinobacteria | Bifidobacteriales | Bifidobacteriaceae | Bifidobacterium |
| Bilophila | Bacteria | Proteobacteria | Deltaproteobacteria | Desulfovibrionales | Desulfovibrionaceae | Bilophila |
| Blautia | Bacteria | Firmicutes | Clostridia | Clostridiales | Lachnospiraceae | Blautia |
| Butyricimonas | Bacteria | Bacteroidetes | Bacteroidia | Bacteroidales | [Odoribacteraceae] | Butyricimonas |
| cc_115 | Bacteria | Firmicutes | Erysipelotrichi | Erysipelotrichales | Erysipelotrichaceae | cc_115 |
| Collinsella | Bacteria | Actinobacteria | Coriobacteriia | Coriobacteriales | Coriobacteriaceae | Collinsella |
| Coprobacillus | Bacteria | Firmicutes | Erysipelotrichi | Erysipelotrichales | Erysipelotrichaceae | Coprobacillus |
| Coprococcus | Bacteria | Firmicutes | Clostridia | Clostridiales | Lachnospiraceae | Coprococcus |
| Eggerthella | Bacteria | Actinobacteria | Coriobacteriia | Coriobacteriales | Coriobacteriaceae | Eggerthella |
| Faecalibacterium | Bacteria | Firmicutes | Clostridia | Clostridiales | Ruminococcaceae | Faecalibacterium |
| Holdemania | Bacteria | Firmicutes | Erysipelotrichi | Erysipelotrichales | Erysipelotrichaceae | Holdemania |
| Lactonifactor | Bacteria | Firmicutes | Clostridia | Clostridiales | Lachnospiraceae | Lactonifactor |
| Odoribacter | Bacteria | Bacteroidetes | Bacteroidia | Bacteroidales | [Odoribacteraceae] | Odoribacter |
| Oscillospira | Bacteria | Firmicutes | Clostridia | Clostridiales | Ruminococcaceae | Oscillospira |
| Parabacteroides | Bacteria | Bacteroidetes | Bacteroidia | Bacteroidales | Porphyromonadaceae | Parabacteroides |
| Phascolarctobacterium | Bacteria | Firmicutes | Clostridia | Clostridiales | Veillonellaceae | Phascolarctobacterium |
| Streptococcus | Bacteria | Firmicutes | Bacilli | Lactobacillales | Streptococcaceae | Streptococcus |
| Sutterella | Bacteria | Proteobacteria | Betaproteobacteria | Burkholderiales | Alcaligenaceae | Sutterella |
| Turicibacter | Bacteria | Firmicutes | Bacilli | Turicibacterales | Turicibacteraceae | Turicibacter |

### Supplementary Table 5. Core Microbiome Comparisons between the Healthy and High-Risk (Depression and Stress Risk) Groups (Study 1)

| Microbiome | Healthy Group | | Mental Health Risk Group | | | | | | | | | |
|---|---|---|---|---|---|---|---|---|---|---|---|---|
| | | | Parenting Stress Cutoff (High) (N = 65) | | Mann-Whitney U test (Stress risk vs Healthy) | | | Depression Cutoff (High) (N = 47) | | Mann-Whitney U test (Depression risk vs Healthy) | | |
| | *Mean* | *SD* | *Mean* | *SD* | *W* | *p*-value | *q*-value | *Mean* | *SD* | *W* | *p*-value | *q*-value |
| Actinomyces | -4.05 | 0.82 | -3.90 | 0.81 | 7393.00 | 0.181 | 0.112 | -3.95 | 0.86 | 5635.00 | 0.529 | 0.489 |
| Bifidobacterium | -1.23 | 0.31 | -1.23 | 0.61 | 7168.00 | 0.101 | **0.076** † | -1.34 | 0.70 | 5956.00 | 0.981 | 0.613 |
| Collinsella | -2.73 | 1.59 | -2.98 | 1.69 | 8563.00 | 0.636 | 0.185 | -2.71 | 1.62 | 5694.50 | 0.611 | 0.514 |
| Eggerthella | -2.94 | 0.76 | -2.87 | 0.77 | 7611.50 | 0.332 | 0.135 | -2.91 | 0.81 | 5668.00 | 0.583 | 0.500 |
| Bacteroides | -0.56 | 0.15 | -0.57 | 0.15 | 9056.00 | 0.227 | 0.116 | -0.58 | 0.20 | 6009.00 | 0.942 | 0.606 |
| Odoribacter | -3.05 | 1.12 | -3.50 | 1.31 | 9733.00 | **0.025** * | **0.024** * | -3.18 | 1.17 | 6529.00 | 0.304 | 0.440 |
| Alistipes | -2.11 | 1.12 | -2.63 | 1.45 | 9880.00 | **0.014** * | **0.019** * | -2.37 | 1.28 | 6616.00 | 0.238 | 0.434 |
| Parabacteroides | -1.87 | 1.06 | -1.95 | 1.24 | 7869.00 | 0.561 | 0.181 | -1.82 | 1.15 | 5148.00 | 0.134 | 0.293 |
| Erysipelatoclostridium | -2.83 | 0.82 | -2.66 | 0.72 | 7046.50 | 0.069 | 0.057 † | -2.72 | 0.68 | 5630.00 | 0.536 | 0.489 |
| Turicibacter | -3.90 | 1.03 | -3.95 | 1.12 | 8531.50 | 0.664 | 0.187 | -3.99 | 1.08 | 6285.50 | 0.547 | 0.489 |
| Streptococcus | -2.09 | 0.66 | -2.03 | 0.72 | 7643.50 | 0.357 | 0.138 | -2.16 | 0.86 | 5908.00 | 0.911 | 0.606 |
| Intestinibacter | -3.45 | 1.04 | -3.32 | 1.01 | 7760.00 | 0.452 | 0.154 | -3.46 | 1.09 | 5913.00 | 0.918 | 0.606 |
| Romboutsia | -3.69 | 1.11 | -3.71 | 1.18 | 8120.00 | 0.834 | 0.217 | -3.86 | 1.22 | 6319.00 | 0.508 | 0.489 |
| Agathobacter | -3.07 | 1.56 | -3.14 | 1.54 | 8568.00 | 0.628 | 0.185 | -3.24 | 1.54 | 6407.00 | 0.411 | 0.462 |
| Anaerostipes | -2.03 | 0.68 | -2.20 | 0.93 | 8974.00 | 0.279 | 0.125 | -2.25 | 0.94 | 6697.50 | 0.184 | 0.384 |
| Blautia | -1.31 | 0.22 | -1.32 | 0.24 | 8501.00 | 0.711 | 0.194 | -1.27 | 0.23 | 5401.00 | 0.300 | 0.440 |
| Dorea | -2.37 | 1.02 | -2.46 | 1.11 | 8590.00 | 0.613 | 0.185 | -2.49 | 1.11 | 6321.00 | 0.520 | 0.489 |
| Fusicatenibacter | -2.37 | 1.23 | -2.81 | 1.53 | 9121.00 | 0.190 | 0.112 | -2.57 | 1.41 | 6226.00 | 0.638 | 0.517 |
| Lachnoclostridium | -1.87 | 0.36 | -1.81 | 0.37 | 7475.00 | 0.240 | 0.118 | -1.75 | 0.34 | 4847.00 | 0.041 * | 0.128 |
| Lachnospira | -2.59 | 1.10 | -2.95 | 1.25 | 9824.00 | **0.018** * | **0.021** * | -3.09 | 1.32 | 7325.00 | **0.013** * | **0.047** * |
| Roseburia | -3.17 | 1.22 | -3.37 | 1.30 | 9088.00 | 0.205 | 0.112 | -3.39 | 1.40 | 6463.00 | 0.362 | 0.440 |
| Sellimonas | -2.98 | 1.13 | -3.01 | 1.12 | 8446.00 | 0.773 | 0.206 | -3.08 | 1.11 | 6385.00 | 0.446 | 0.476 |
| Tuzzerella | -3.87 | 1.05 | -3.98 | 1.06 | 8725.00 | 0.459 | 0.154 | -4.04 | 1.08 | 6432.50 | 0.376 | 0.445 |
| Lachnospiraceae_UCG-004 | -3.59 | 1.19 | -3.69 | 1.22 | 8572.50 | 0.620 | 0.185 | -3.55 | 1.24 | 5865.50 | 0.845 | 0.578 |
| Colidextribacter | -3.02 | 1.05 | -3.21 | 1.12 | 8950.00 | 0.293 | 0.127 | -3.02 | 1.05 | 5926.50 | 0.938 | 0.606 |
| Flavonifractor | -2.95 | 0.76 | -2.90 | 0.67 | 8244.00 | 0.987 | 0.246 | -2.95 | 0.81 | 5750.00 | 0.689 | 0.529 |
| Oscillibacter | -2.85 | 0.89 | -3.03 | 1.03 | 8992.00 | 0.266 | 0.122 | -2.92 | 0.96 | 6188.50 | 0.689 | 0.529 |
| Butyricicoccus | -2.56 | 0.53 | -2.74 | 0.75 | 9141.00 | 0.182 | 0.112 | -2.66 | 0.63 | 6202.00 | 0.671 | 0.529 |
| Faecalibacterium | -1.46 | 0.88 | -1.79 | 1.29 | 9364.00 | 0.095 | 0.075 † | -1.88 | 1.38 | 7038.50 | 0.051 | 0.131 |
| Negativibacillus | -3.90 | 1.05 | -4.12 | 1.04 | 9143.50 | 0.158 | 0.109 | -4.00 | 1.02 | 6304.50 | 0.521 | 0.489 |
| Subdoligranulum | -3.02 | 1.33 | -3.26 | 1.46 | 8812.00 | 0.396 | 0.144 | -3.33 | 1.46 | 6594.00 | 0.249 | 0.436 |
| UBA1819 | -3.66 | 0.90 | -3.85 | 1.04 | 9170.00 | 0.163 | 0.109 | -3.84 | 1.05 | 6489.50 | 0.337 | 0.440 |
| Monoglobus | -2.84 | 1.13 | -3.28 | 1.21 | 10195.00 | **0.003** ** | **0.005** ** | -3.08 | 1.18 | 6633.00 | 0.224 | 0.426 |
| Phascolarctobacterium | -3.10 | 1.46 | -3.63 | 1.52 | 9823.00 | **0.015** * | **0.019** * | -3.30 | 1.59 | 6129.00 | 0.765 | 0.555 |
| Veillonella | -3.30 | 1.49 | -2.89 | 1.57 | 6803.00 | **0.025** * | **0.024** * | -3.20 | 1.58 | 5676.00 | 0.582 | 0.500 |
| Bilophila | -3.04 | 1.13 | -3.31 | 1.26 | 9023.00 | 0.243 | 0.118 | -3.19 | 1.25 | 6132.00 | 0.765 | 0.555 |
| Parasutterella | -3.11 | 1.43 | -3.40 | 1.61 | 8920.00 | 0.306 | 0.127 | -3.37 | 1.52 | 6505.00 | 0.319 | 0.440 |
| Sutterella | -2.94 | 1.62 | -3.53 | 1.64 | 9882.00 | **0.011** * | **0.016** * | -3.25 | 1.69 | 6459.00 | 0.357 | 0.440 |
| Escherichia-Shigella | -3.60 | 1.32 | -3.18 | 1.34 | 6747.00 | **0.019** * | **0.021** * | -3.26 | 1.40 | 5055.00 | 0.084 | 0.204 |

**$p$ or $q$ <.01, *$p$ or $q$ <.05, †$q$ <.10.

### Supplementary Table 6. Statistical Comparisons between the Healthy and High- and Low-Resilience Groups (Study 1)

| | | Healthy Group (N = 254) | | High Resilience Group (N = 35) | | Low Resilience Group (N = 30) | | Kruskal-Wallis test ( Healthy vs High resi vs Low resi) | | | | |
|---|---|---|---|---|---|---|---|---|---|---|---|---|
| | | *Mean* | *SD* | *Mean* | *SD* | *Mean* | *SD* | *t* | *p* -value | | *q* -value | |
| Psychological and physical indices | Parenting Stress (PSI)_total | 165.19 | 27.08 | 215.97 | 13.37 | 231.11 | 22.75 | 131.40 | **0.000** | ** | **0.000** | ** |
| | Parenting Stress (PSI)_child | 72.47 | 14.57 | 89.94 | 13.84 | 94.61 | 15.13 | 62.17 | **0.000** | ** | **0.000** | ** |
| | Parenting Stress (PSI)_parent | 92.73 | 16.23 | 126.03 | 9.89 | 136.50 | 11.28 | 134.10 | **0.000** | ** | **0.000** | ** |
| | Depression (BDI) | 7.89 | 4.82 | 12.60 | 5.15 | 27.29 | 6.82 | 94.54 | **0.000** | ** | **0.000** | ** |
| | Physical activity index (MDPS_PAI) | 2.71 | 1.44 | 3.86 | 1.29 | 4.63 | 1.00 | 56.28 | **0.000** | ** | **0.000** | ** |
| | Somatic disorders index (MDPS_SDI) | 0.78 | 1.04 | 1.40 | 1.46 | 1.83 | 1.53 | 20.65 | **0.000** | ** | **0.000** | ** |
| | Hormone activity index (MDPS_HAI) | 3.96 | 1.72 | 4.66 | 1.49 | 5.63 | 1.50 | 25.03 | **0.000** | ** | **0.000** | ** |
| | Microvascular disorders index (MDPS_MDI) | 3.40 | 1.94 | 4.00 | 1.88 | 5.07 | 1.84 | 19.01 | **0.000** | ** | **0.000** | ** |
| | Meteoropathy related index (MDPS_MRI) | 2.50 | 1.56 | 2.69 | 1.76 | 3.93 | 1.53 | 18.46 | **0.000** | ** | **0.000** | ** |
| | sleeping time (hours) | 6.80 | 1.07 | 6.74 | 1.07 | 6.50 | 1.08 | 2.29 | 0.319 | | 0.194 | |
| | sleeping quality | 1.98 | 0.59 | 1.89 | 0.47 | 1.60 | 0.50 | 12.07 | **0.002** | ** | **0.005** | ** |
| Dietary intake | staple food | 4.55 | 0.90 | 4.54 | 0.89 | 4.40 | 0.89 | 2.31 | 0.315 | | 0.194 | |
| | unrefined grain | 1.44 | 0.98 | 1.31 | 0.72 | 1.33 | 0.99 | 1.08 | 0.588 | | 0.252 | |
| | root vegetable | 3.32 | 1.06 | 3.06 | 0.94 | 3.40 | 1.19 | 3.03 | 0.221 | | 0.159 | |
| | green and yellow vegetables | 3.06 | 1.02 | 3.11 | 1.11 | 3.23 | 1.17 | 0.42 | 0.811 | | 0.295 | |
| | light colored vegetables | 3.19 | 1.01 | 3.23 | 1.00 | 3.37 | 1.07 | 0.78 | 0.681 | | 0.266 | |
| | fruits | 2.65 | 1.06 | 2.66 | 1.03 | 2.57 | 1.19 | 0.40 | 0.819 | | 0.295 | |
| | meat | 3.56 | 0.93 | 3.37 | 1.09 | 3.27 | 1.05 | 2.87 | 0.239 | | 0.163 | |
| | fish and shellfish | 2.62 | 0.84 | 2.57 | 0.85 | 2.57 | 1.01 | 0.78 | 0.677 | | 0.266 | |
| | egg | 3.04 | 0.85 | 2.86 | 0.97 | 3.03 | 1.00 | 0.63 | 0.735 | | 0.283 | |
| | milk and cheese | 3.05 | 1.17 | 2.83 | 1.07 | 2.60 | 1.30 | 4.27 | 0.119 | | 0.106 | |
| | yogurt and lactic fermenting beverage | 2.31 | 1.07 | 2.43 | 1.14 | 2.00 | 0.95 | 2.14 | 0.344 | | 0.198 | |
| | soy product | 2.63 | 0.90 | 2.34 | 0.87 | 2.63 | 1.00 | 3.29 | 0.194 | | 0.144 | |
| | natto | 1.84 | 0.85 | 2.00 | 0.87 | 2.00 | 1.05 | 1.37 | 0.506 | | 0.229 | |
| | pickled vegetables | 1.73 | 0.75 | 1.66 | 0.76 | 1.63 | 0.89 | 1.81 | 0.407 | | 0.209 | |
| | seaweed | 2.10 | 0.74 | 2.14 | 0.81 | 2.23 | 1.04 | 0.10 | 0.950 | | 0.313 | |
| | mushroom | 2.60 | 0.85 | 2.51 | 0.98 | 2.73 | 0.87 | 1.22 | 0.545 | | 0.242 | |
| | snack | 3.50 | 0.97 | 3.37 | 0.77 | 3.37 | 1.27 | 0.96 | 0.621 | | 0.256 | |
| | sugary drink | 2.89 | 1.19 | 2.77 | 1.09 | 3.07 | 1.26 | 1.02 | 0.602 | | 0.252 | |
| | Japanese Diet Index | 17.12 | 4.20 | 16.94 | 4.27 | 17.67 | 5.56 | 0.35 | 0.843 | | 0.299 | |
| Microbiome | Shannon α | 6.16 | 0.47 | 6.00 | 0.53 | 5.95 | 0.54 | 5.04 | 0.080 | | 0.086 | † |
| | Actinomyces | -4.05 | 0.82 | -3.93 | 0.77 | -3.87 | 0.88 | 1.93 | 0.384 | | 0.206 | |
| | Bifidobacterium | -1.23 | 0.31 | -1.17 | 0.29 | -1.31 | 0.84 | 2.79 | 0.248 | | 0.163 | |
| | Collinsella | -2.73 | 1.59 | -3.00 | 1.69 | -2.96 | 1.72 | 0.25 | 0.882 | | 0.299 | |
| | Eggerthella | -2.94 | 0.76 | -3.06 | 0.91 | -2.64 | 0.48 | 4.54 | 0.103 | | 0.099 | † |
| | Bacteroides | -0.56 | 0.15 | -0.59 | 0.14 | -0.55 | 0.16 | 3.65 | 0.162 | | 0.129 | |
| | Odoribacter | -3.05 | 1.12 | -3.68 | 1.34 | -3.29 | 1.26 | 6.25 | **0.043** | * | **0.060** | † |
| | Alistipes | -2.11 | 1.12 | -2.69 | 1.50 | -2.56 | 1.42 | 6.22 | **0.044** | * | **0.060** | † |
| | Parabacteroides | -1.87 | 1.06 | -2.01 | 1.21 | -1.87 | 1.28 | 3.83 | 0.148 | | 0.126 | |
| | Erysipelatoclostridium | -2.83 | 0.82 | -2.71 | 0.78 | -2.60 | 0.65 | 3.41 | 0.182 | | 0.140 | |
| | Turicibacter | -3.90 | 1.03 | -3.91 | 1.14 | -3.99 | 1.12 | 0.26 | 0.881 | | 0.299 | |
| | Streptococcus | -2.09 | 0.66 | -1.99 | 0.61 | -2.07 | 0.84 | 0.85 | 0.655 | | 0.265 | |
| | Intestinibacter | -3.45 | 1.04 | -3.32 | 0.99 | -3.32 | 1.05 | 0.57 | 0.754 | | 0.285 | |
| | Romboutsia | -3.69 | 1.11 | -3.65 | 1.18 | -3.77 | 1.20 | 0.23 | 0.894 | | 0.299 | |
| | Agathobacter | -3.07 | 1.56 | -2.75 | 1.43 | -3.59 | 1.56 | 4.99 | 0.082 | | 0.086 | † |
| | Anaerostipes | -2.03 | 0.68 | -2.10 | 0.84 | -2.31 | 1.03 | 1.74 | 0.421 | | 0.211 | |
| | Blautia | -1.31 | 0.22 | -1.37 | 0.25 | -1.26 | 0.22 | 4.26 | 0.118 | | 0.106 | |
| | Dorea | -2.37 | 1.02 | -2.34 | 1.07 | -2.60 | 1.15 | 1.37 | 0.506 | | 0.229 | |
| | Fusicatenibacter | -2.37 | 1.23 | -2.81 | 1.48 | -2.82 | 1.60 | 2.15 | 0.343 | | 0.198 | |
| | Lachnoclostridium | -1.87 | 0.36 | -1.90 | 0.35 | -1.70 | 0.36 | 6.23 | **0.044** | * | **0.060** | † |
| | Lachnospira | -2.59 | 1.10 | -2.88 | 1.24 | -3.02 | 1.28 | 5.87 | 0.052 | | 0.067 | † |
| | Roseburia | -3.17 | 1.22 | -3.20 | 1.14 | -3.58 | 1.45 | 2.05 | 0.360 | | 0.203 | |
| | Sellimonas | -2.98 | 1.13 | -3.06 | 1.12 | -2.94 | 1.13 | 0.41 | 0.817 | | 0.295 | |
| | Tuzzerella | -3.87 | 1.05 | -3.86 | 1.05 | -4.13 | 1.06 | 1.44 | 0.488 | | 0.229 | |
| | Lachnospiraceae_UCG-004 | -3.59 | 1.19 | -3.70 | 1.20 | -3.69 | 1.27 | 0.26 | 0.880 | | 0.299 | |
| | Colidextribacter | -3.02 | 1.05 | -3.28 | 1.11 | -3.12 | 1.14 | 2.52 | 0.286 | | 0.183 | |
| | Flavonifractor | -2.95 | 0.76 | -3.02 | 0.63 | -2.76 | 0.69 | 6.72 | **0.034** | * | **0.060** | † |
| | Oscillibacter | -2.85 | 0.89 | -3.05 | 0.99 | -3.01 | 1.08 | 1.48 | 0.479 | | 0.229 | |
| | Butyricicoccus | -2.56 | 0.53 | -2.77 | 0.89 | -2.71 | 0.58 | 1.86 | 0.396 | | 0.208 | |
| | Faecalibacterium | -1.46 | 0.88 | -1.54 | 0.96 | -2.09 | 1.55 | 3.69 | 0.159 | | 0.129 | |
| | Negativibacillus | -3.90 | 1.05 | -4.14 | 1.05 | -4.10 | 1.06 | 2.00 | 0.370 | | 0.203 | |
| | Subdoligranulum | -3.02 | 1.33 | -3.13 | 1.47 | -3.42 | 1.46 | 1.50 | 0.475 | | 0.229 | |
| | UBA1819 | -3.66 | 0.90 | -4.03 | 1.05 | -3.64 | 1.02 | 4.59 | 0.101 | | 0.099 | † |
| | Monoglobus | -2.84 | 1.13 | -3.16 | 1.17 | -3.42 | 1.27 | 8.85 | **0.011** | * | **0.023** | * |
| | Phascolarctobacterium | -3.10 | 1.46 | -3.81 | 1.42 | -3.43 | 1.63 | 8.56 | **0.013** | * | **0.025** | * |
| | Veillonella | -3.30 | 1.49 | -2.89 | 1.58 | -2.90 | 1.57 | 5.05 | 0.079 | | 0.086 | † |
| | Bilophila | -3.04 | 1.13 | -3.42 | 1.27 | -3.17 | 1.27 | 2.84 | 0.243 | | 0.163 | |
| | Parasutterella | -3.11 | 1.43 | -3.37 | 1.64 | -3.42 | 1.60 | 1.07 | 0.590 | | 0.252 | |
| | Sutterella | -2.94 | 1.62 | -3.56 | 1.63 | -3.49 | 1.68 | 6.48 | **0.038** | * | **0.060** | † |
| | Escherichia-Shigella | -3.60 | 1.32 | -3.25 | 1.32 | -3.09 | 1.38 | 5.76 | 0.055 | | 0.067 | † |

**$p$ or $q$ <.01, *$p$ or $q$ <.05, †$q$ <.10.

### Supplementary Table 7. Statistical Comparisons between High and Low JDI12 Groups (Study 1)

| | | High JDI Group (N = 173) | | Low JDI Group (N = 174) | | welch's t-test (High JDI vs Low JDI) | | | | | |
|---|---|---|---|---|---|---|---|---|---|---|---|
| | | *Mean* | | *Mean* | | *t* | | *p-value* | | *q-value* | |
| | Parenting Stress (PSI)_total | 174.76 | 33.74 | 178.29 | 34.70 | -0.93 | 321.47 | 0.354 | | 0.533 | |
| | Parenting Stress (PSI)_child | 75.26 | 16.20 | 77.09 | 16.77 | -1.00 | 321.30 | 0.319 | | 0.519 | |
| | Parenting Stress (PSI)_parent | 99.50 | 20.90 | 101.20 | 21.84 | -0.72 | 320.98 | 0.474 | | 0.551 | |
| | Depression (BDI) | 10.08 | 7.45 | 11.93 | 8.73 | -2.09 | 329.03 | **0.037** | * | **0.086** | † |
| Psychological and physical indices | Physical activity index (MDPS_PAI) | 2.88 | 1.45 | 3.25 | 1.59 | -2.26 | 342.33 | **0.025** | * | **0.059** | † |
| | Somatic disorders index (MDPS_SDI) | 0.91 | 1.21 | 1.03 | 1.24 | -0.92 | 344.87 | 0.356 | | 0.533 | |
| | Hormone activity index (MDPS_HAI) | 4.10 | 1.78 | 4.35 | 1.72 | -1.31 | 344.56 | 0.190 | | 0.382 | |
| | Microvascular disorders index (MDPS_MDI) | 3.38 | 1.90 | 3.87 | 2.07 | -2.31 | 342.94 | **0.022** | * | **0.057** | † |
| | Meteoropathy related index (MDPS_MRI) | 2.54 | 1.59 | 2.87 | 1.71 | -1.86 | 343.30 | 0.063 | | 0.140 | |
| | sleeping time (hours) | 6.77 | 1.07 | 6.76 | 1.11 | 0.05 | 343.44 | 0.961 | | 0.775 | |
| | sleeping quality | 1.93 | 0.58 | 1.90 | 0.62 | 0.53 | 343.59 | 0.596 | | 0.63 | |
| | staple food | 4.62 | 0.84 | 4.39 | 0.98 | 2.45 | 337.64 | **0.015** | * | **0.046** | * |
| | unrefined grain | 1.66 | 1.13 | 1.14 | 0.60 | 5.30 | 260.01 | **0.000** | ** | **0.000** | ** |
| | root vegetable | 3.77 | 0.92 | 2.80 | 0.95 | 9.63 | 344.75 | **0.000** | ** | **0.000** | ** |
| | green and yellow vegetables | 3.64 | 0.96 | 2.49 | 0.78 | 12.29 | 330.81 | **0.000** | ** | **0.000** | ** |
| | light colored vegetables | 3.68 | 0.96 | 2.75 | 0.87 | 9.46 | 341.44 | **0.000** | ** | **0.000** | ** |
| | fruits | 3.23 | 0.94 | 2.05 | 0.83 | 12.29 | 339.72 | **0.000** | ** | **0.000** | ** |
| | meat | 3.77 | 0.89 | 3.24 | 0.97 | 5.36 | 342.74 | **0.000** | ** | **0.000** | ** |
| | fish and shellfish | 2.99 | 0.83 | 2.22 | 0.69 | 9.47 | 332.45 | **0.000** | ** | **0.000** | ** |
| | egg | 3.26 | 0.85 | 2.76 | 0.84 | 5.54 | 344.93 | **0.000** | ** | **0.000** | ** |
| Dietary intake | milk and cheese | 3.27 | 1.12 | 2.67 | 1.15 | 4.95 | 344.81 | **0.000** | ** | **0.000** | ** |
| | yogurt and lactic fermenting beverage | 2.68 | 1.13 | 1.92 | 0.88 | 6.99 | 324.60 | **0.000** | ** | **0.000** | ** |
| | soy product | 3.10 | 0.92 | 2.09 | 0.62 | 11.99 | 301.03 | **0.000** | ** | **0.000** | ** |
| | natto | 2.29 | 0.93 | 1.52 | 0.65 | 8.91 | 308.58 | **0.000** | ** | **0.000** | ** |
| | pickled vegetables | 1.84 | 0.87 | 1.54 | 0.58 | 3.83 | 297.60 | **0.000** | ** | **0.001** | ** |
| | seaweed | 2.45 | 0.77 | 1.75 | 0.58 | 9.43 | 319.33 | **0.000** | ** | **0.000** | ** |
| | mushroom | 3.04 | 0.81 | 2.10 | 0.64 | 11.98 | 326.03 | **0.000** | ** | **0.000** | ** |
| | snack | 3.57 | 0.96 | 3.32 | 1.05 | 2.32 | 342.39 | **0.021** | * | **0.057** | † |
| | sugary drink | 2.71 | 1.15 | 3.05 | 1.23 | -2.63 | 343.53 | **0.009** | ** | **0.029** | * |
| | Japanese Diet Index | 20.47 | 3.18 | 13.68 | 1.93 | 24.04 | 282.77 | **0.000** | ** | **0.000** | ** |
| | Shannonα | 6.15 | 0.47 | 6.08 | 0.49 | 1.35 | 344.11 | 0.179 | | 0.381 | |
| | Actinomyces | -4.04 | 0.81 | -4.00 | 0.83 | -0.47 | 344.81 | 0.638 | | 0.631 | |
| | Bifidobacterium | -1.23 | 0.33 | -1.24 | 0.43 | 0.14 | 324.20 | 0.887 | | 0.767 | |
| | Collinsella | -2.79 | 1.58 | -2.73 | 1.62 | -0.34 | 344.84 | 0.733 | | 0.677 | |
| | Eggerthella | -2.92 | 0.76 | -2.96 | 0.79 | 0.48 | 344.51 | 0.631 | | 0.631 | |
| | Bacteroides | -0.56 | 0.15 | -0.56 | 0.16 | -0.16 | 341.98 | 0.874 | | 0.767 | |
| | Odoribacter | -3.14 | 1.15 | -3.12 | 1.17 | -0.16 | 344.96 | 0.875 | | 0.767 | |
| | Alistipes | -2.18 | 1.22 | -2.25 | 1.18 | 0.51 | 344.38 | 0.610 | | 0.631 | |
| | Parabacteroides | -1.82 | 0.98 | -1.94 | 1.18 | 1.07 | 333.63 | 0.286 | | 0.481 | |
| | Erysipelatoclostridium | -2.82 | 0.80 | -2.77 | 0.78 | -0.57 | 344.73 | 0.571 | | 0.631 | |
| | Turicibacter | -3.94 | 1.03 | -3.88 | 1.06 | -0.49 | 344.92 | 0.626 | | 0.631 | |
| | Streptococcus | -2.07 | 0.62 | -2.12 | 0.75 | 0.71 | 334.42 | 0.477 | | 0.551 | |
| Microbiome | Intestinibacter | -3.41 | 1.04 | -3.46 | 1.05 | 0.48 | 344.98 | 0.629 | | 0.631 | |
| | Romboutsia | -3.76 | 1.13 | -3.66 | 1.13 | -0.85 | 344.93 | 0.399 | | 0.551 | |
| | Agathobacter | -2.86 | 1.48 | -3.23 | 1.59 | 2.26 | 343.44 | **0.025** | * | **0.059** | † |
| | Anaerostipes | -2.07 | 0.79 | -2.07 | 0.68 | 0.04 | 337.10 | 0.965 | | 0.775 | |
| | Blautia | -1.32 | 0.23 | -1.31 | 0.22 | -0.42 | 343.19 | 0.671 | | 0.652 | |
| | Dorea | -2.33 | 0.99 | -2.46 | 1.10 | 1.13 | 341.64 | 0.257 | | 0.460 | |
| | Fusicatenibacter | -2.43 | 1.30 | -2.48 | 1.31 | 0.35 | 345.00 | 0.723 | | 0.677 | |
| | Lachnoclostridium | -1.85 | 0.35 | -1.87 | 0.37 | 0.74 | 344.26 | 0.458 | | 0.551 | |
| | Lachnospira | -2.63 | 1.17 | -2.77 | 1.16 | 1.07 | 344.96 | 0.287 | | 0.481 | |
| | Roseburia | -3.20 | 1.18 | -3.20 | 1.29 | -0.02 | 342.66 | 0.981 | | 0.777 | |
| | Sellimonas | -3.04 | 1.13 | -2.95 | 1.10 | -0.74 | 344.70 | 0.459 | | 0.551 | |
| | Tuzzerella | -3.82 | 1.05 | -3.96 | 1.06 | 1.26 | 344.99 | 0.209 | | 0.394 | |
| | Lachnospiraceae_UCG-004 | -3.55 | 1.19 | -3.63 | 1.22 | 0.59 | 344.83 | 0.558 | | 0.631 | |
| | Colidextribacter | -3.06 | 1.08 | -3.03 | 1.03 | -0.31 | 344.03 | 0.759 | | 0.689 | |
| | Flavonifractor | -2.92 | 0.73 | -2.98 | 0.76 | 0.76 | 344.64 | 0.449 | | 0.551 | |
| | Oscillibacter | -2.87 | 0.95 | -2.91 | 0.89 | 0.40 | 342.97 | 0.691 | | 0.660 | |
| | Butyricicoccus | -2.57 | 0.53 | -2.63 | 0.68 | 0.89 | 326.58 | 0.375 | | 0.546 | |
| | Faecalibacterium | -1.52 | 0.97 | -1.53 | 0.97 | 0.10 | 344.99 | 0.922 | | 0.774 | |
| | Negativibacillus | -3.90 | 1.05 | -3.99 | 1.04 | 0.74 | 344.93 | 0.459 | | 0.551 | |
| | Subdoligranulum | -2.90 | 1.29 | -3.24 | 1.42 | 2.35 | 342.29 | **0.019** | * | **0.056** | † |
| | UBA1819 | -3.69 | 0.96 | -3.77 | 0.92 | 0.73 | 344.29 | 0.465 | | 0.551 | |
| | Monoglobus | -2.91 | 1.18 | -2.92 | 1.12 | 0.12 | 343.70 | 0.903 | | 0.769 | |
| | Phascolarctobacterium | -3.13 | 1.47 | -3.28 | 1.49 | 0.94 | 344.97 | 0.347 | | 0.533 | |
| | Veillonella | -3.30 | 1.52 | -3.18 | 1.52 | -0.76 | 345.00 | 0.448 | | 0.551 | |
| | Bilophila | -3.03 | 1.14 | -3.19 | 1.19 | 1.25 | 344.70 | 0.214 | | 0.394 | |
| | Parasutterella | -3.13 | 1.45 | -3.25 | 1.49 | 0.79 | 344.84 | 0.431 | | 0.551 | |
| | Sutterella | -3.03 | 1.63 | -3.02 | 1.64 | -0.06 | 345.00 | 0.949 | | 0.775 | |
| | Escherichia-Shigella | -3.62 | 1.34 | -3.43 | 1.32 | -1.30 | 344.85 | 0.193 | | 0.382 | |

**$p$ or $q$ <.01, *$p$ or $q$ <.05, †$q$ <.10.

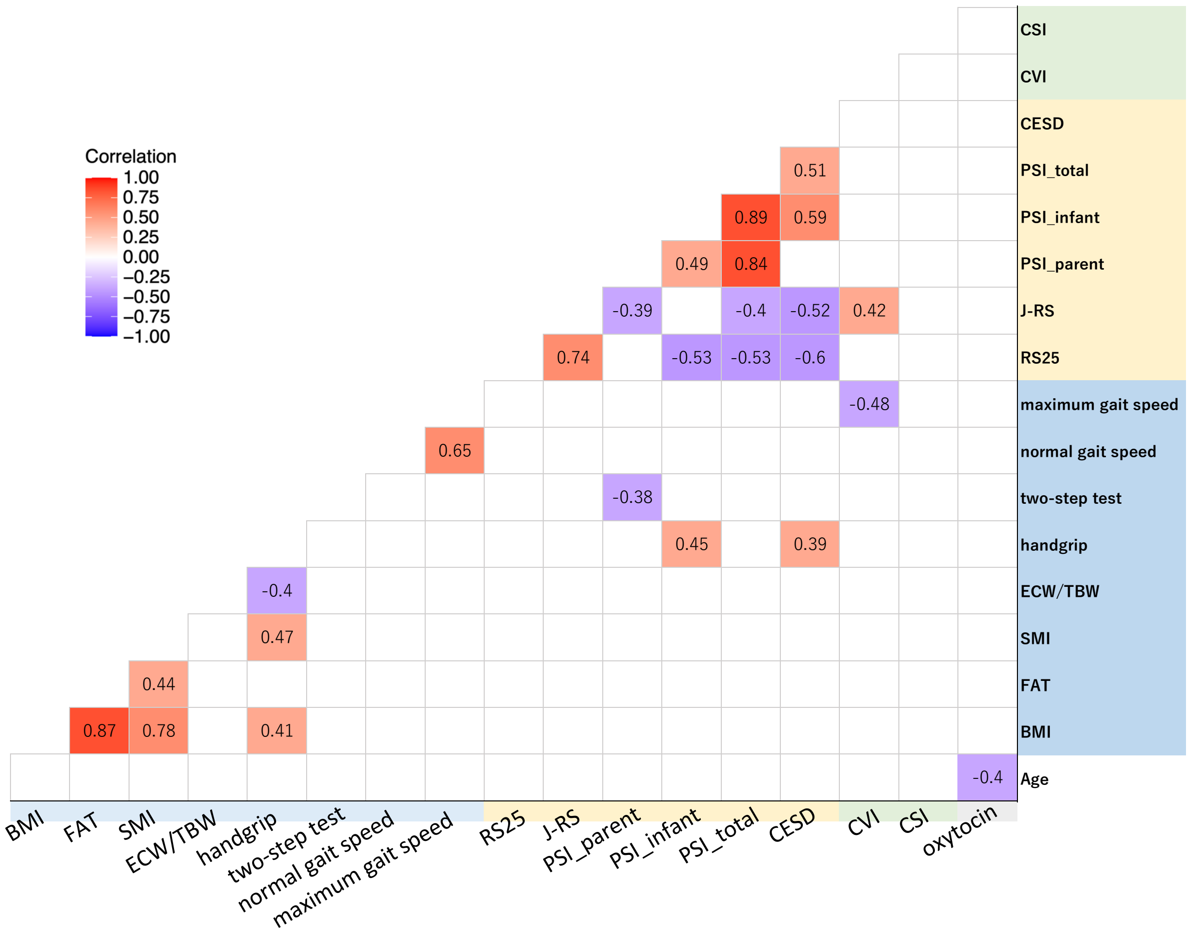


**Supplementary Fig. 1. Pearson's correlations between physical function, physiological indices, and psychological indices in postpartum women (Study 2).**

Values in the figure represent *r* at p <.05.

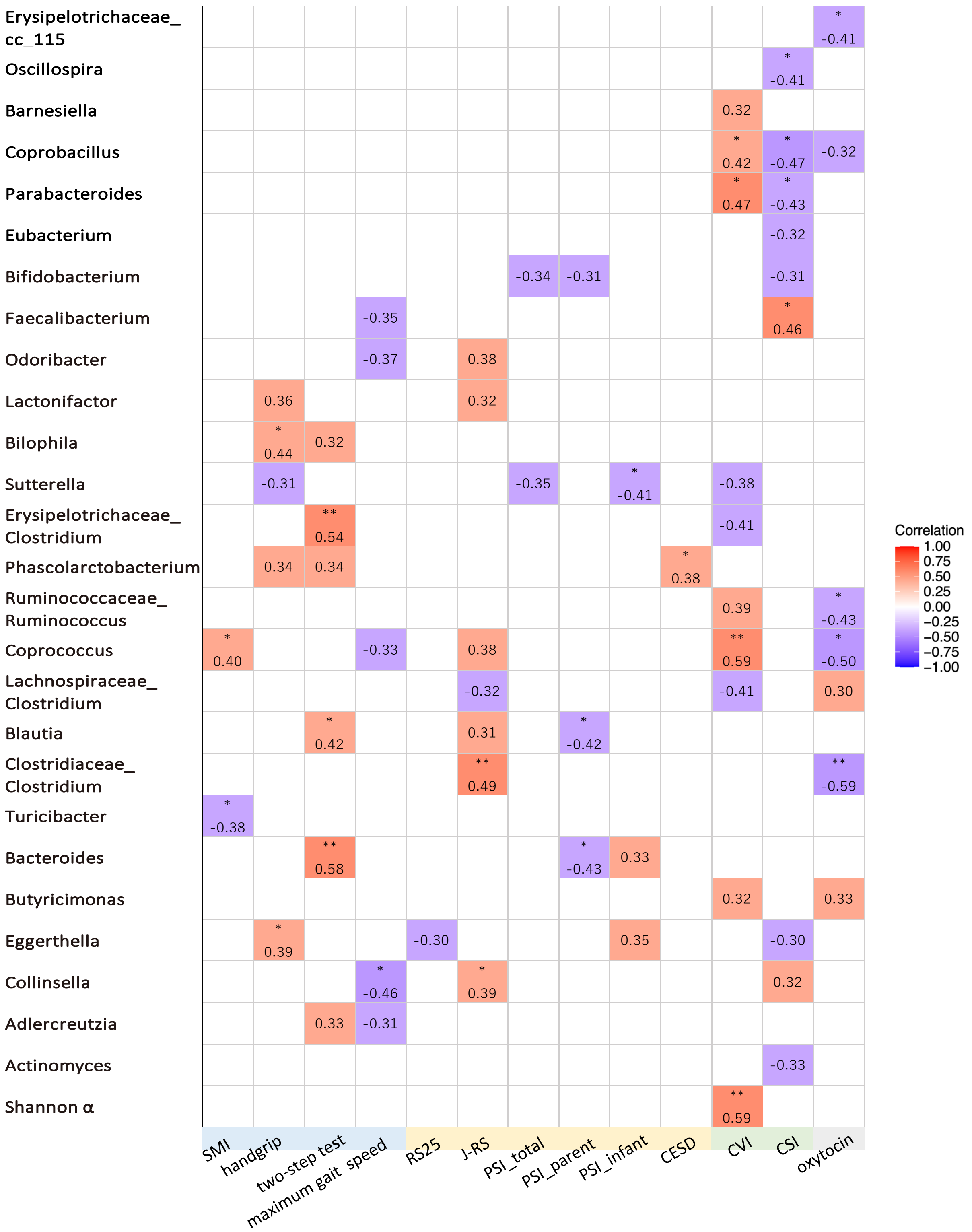


**Supplementary Fig. 2. Pearson's correlations between intestinal microbiota and physical and psychological indices (Study 2).**

Values in the figure represent *r* with significant or medium to large effect size. **$p$ <.01, *$p$ <.05.

**Supplementary Fig. 3. Identifying the intestinal microbiome by shotgun metagenomic analysis (Study 2).**

| Clostridium | Subject s1 | s2 | s3 | s4 | s5 | s6 | s7 | s8 | s9 | s10 | s11 | s12 | s13 | s14 | s15 | s16 | s17 | s18 | s19 |
|---|---|---|---|---|---|---|---|---|---|---|---|---|---|---|---|---|---|---|---|
| **Clostridium sp. SY8519** | **0.00082** | **0.00029** | **0.00069** | **0.00106** | **0.00099** | **0.00038** | **0.00095** | **0.00060** | **0.00020** | **0.00049** | **0.00056** | **0.00066** | **0.00019** | **0.00089** | **0.00036** | **0.00014** | **0.00069** | **0.00016** | **0.00087** |
| Clostridium aceticum | 0.00003 | 0.00001 | 0.00002 | 0.00005 | 0.00002 | 0.00002 | 0.00003 | 0.00003 | 0.00001 | 0.00002 | 0.00001 | 0.00003 | 0.00002 | 0.00005 | 0.00002 | 0.00001 | 0.00003 | 0.00001 | 0.00003 |
| Clostridium acetobutylicum | 0.00002 | 0.00001 | 0.00002 | 0.00002 | 0.00002 | 0.00001 | 0.00001 | 0.00003 | 0.00001 | 0.00001 | 0.00002 | 0.00001 | 0.00001 | 0.00008 | 0.00001 | 0.00001 | 0.00002 | 0.00000 | 0.00002 |
| Clostridium argentinense | 0.00003 | 0.00002 | 0.00001 | 0.00005 | 0.00002 | 0.00002 | 0.00003 | 0.00004 | 0.00001 | 0.00002 | 0.00002 | 0.00002 | 0.00002 | 0.00011 | 0.00002 | 0.00002 | 0.00006 | 0.00001 | 0.00004 |
| Clostridium autoethanogenum | 0.00000 | 0.00000 | 0.00000 | 0.00000 | 0.00000 | 0.00001 | 0.00000 | 0.00000 | 0.00000 | 0.00000 | 0.00000 | 0.00000 | 0.00000 | 0.00001 | 0.00000 | 0.00000 | 0.00000 | 0.00000 | 0.00000 |
| Clostridium baratii | 0.00004 | 0.00002 | 0.00003 | 0.00004 | 0.00002 | 0.00002 | 0.00003 | 0.00004 | 0.00001 | 0.00002 | 0.00002 | 0.00003 | 0.00001 | 0.00016 | 0.00002 | 0.00008 | 0.00003 | 0.00002 | 0.00004 |
| Clostridium beijerinckii | 0.00003 | 0.00003 | 0.00003 | 0.00006 | 0.00004 | 0.00001 | 0.00004 | 0.00006 | 0.00001 | 0.00003 | 0.00004 | 0.00003 | 0.00001 | 0.00012 | 0.00003 | 0.00007 | 0.00005 | 0.00002 | 0.00007 |
| Clostridium bornimense | 0.00005 | 0.00002 | 0.00002 | 0.00007 | 0.00003 | 0.00003 | 0.00004 | 0.00004 | 0.00001 | 0.00003 | 0.00002 | 0.00003 | 0.00001 | 0.00015 | 0.00003 | 0.00003 | 0.00005 | 0.00001 | 0.00004 |
| Clostridium botulinum | 0.00013 | 0.00007 | 0.00010 | 0.00019 | 0.00011 | 0.00005 | 0.00009 | 0.00016 | 0.00007 | 0.00007 | 0.00008 | 0.00011 | 0.00006 | 0.00057 | 0.00009 | 0.00019 | 0.00014 | 0.00003 | 0.00018 |
| Clostridium butyricum | 0.00009 | 0.00005 | 0.00005 | 0.00013 | 0.00006 | 0.00004 | 0.00010 | 0.00049 | 0.00003 | 0.00007 | 0.00007 | 0.00008 | 0.00003 | 0.00022 | 0.00006 | 0.00007 | 0.00009 | 0.00004 | 0.00011 |
| Clostridium carboxidivorans | 0.00002 | 0.00001 | 0.00001 | 0.00003 | 0.00002 | 0.00000 | 0.00003 | 0.00004 | 0.00001 | 0.00001 | 0.00001 | 0.00002 | 0.00001 | 0.00009 | 0.00002 | 0.00002 | 0.00002 | 0.00001 | 0.00003 |
| Clostridium cellulovorans | 0.00004 | 0.00002 | 0.00002 | 0.00005 | 0.00003 | 0.00001 | 0.00004 | 0.00004 | 0.00001 | 0.00003 | 0.00002 | 0.00003 | 0.00001 | 0.00013 | 0.00003 | 0.00003 | 0.00006 | 0.00002 | 0.00007 |
| Clostridium chauvoei | 0.00001 | 0.00001 | 0.00001 | 0.00001 | 0.00002 | 0.00001 | 0.00001 | 0.00002 | 0.00000 | 0.00001 | 0.00001 | 0.00001 | 0.00001 | 0.00010 | 0.00002 | 0.00002 | 0.00002 | 0.00001 | 0.00001 |
| Clostridium cochlearium | 0.00001 | 0.00001 | 0.00001 | 0.00001 | 0.00001 | 0.00000 | 0.00001 | 0.00001 | 0.00000 | 0.00001 | 0.00001 | 0.00001 | 0.00000 | 0.00008 | 0.00001 | 0.00002 | 0.00001 | 0.00001 | 0.00003 |
| Clostridium diolis | 0.00000 | 0.00000 | 0.00000 | 0.00000 | 0.00000 | 0.00000 | 0.00000 | 0.00001 | 0.00000 | 0.00000 | 0.00000 | 0.00000 | 0.00000 | 0.00000 | 0.00000 | 0.00001 | 0.00000 | 0.00000 | 0.00000 |
| Clostridium drakei | 0.00002 | 0.00000 | 0.00001 | 0.00002 | 0.00001 | 0.00001 | 0.00001 | 0.00001 | 0.00000 | 0.00001 | 0.00001 | 0.00001 | 0.00000 | 0.00003 | 0.00001 | 0.00001 | 0.00001 | 0.00001 | 0.00001 |
| Clostridium estertheticum | 0.00003 | 0.00002 | 0.00001 | 0.00006 | 0.00002 | 0.00002 | 0.00003 | 0.00003 | 0.00001 | 0.00002 | 0.00001 | 0.00003 | 0.00000 | 0.00009 | 0.00002 | 0.00004 | 0.00003 | 0.00001 | 0.00004 |
| Clostridium formicaceticum | 0.00004 | 0.00001 | 0.00001 | 0.00003 | 0.00001 | 0.00001 | 0.00002 | 0.00002 | 0.00001 | 0.00001 | 0.00001 | 0.00001 | 0.00001 | 0.00006 | 0.00002 | 0.00001 | 0.00002 | 0.00000 | 0.00002 |
| Clostridium intestinale | 0.00002 | 0.00001 | 0.00001 | 0.00003 | 0.00001 | 0.00001 | 0.00002 | 0.00002 | 0.00001 | 0.00002 | 0.00002 | 0.00002 | 0.00001 | 0.00012 | 0.00002 | 0.00003 | 0.00004 | 0.00001 | 0.00004 |
| Clostridium isatidis | 0.00001 | 0.00002 | 0.00001 | 0.00002 | 0.00001 | 0.00001 | 0.00001 | 0.00003 | 0.00001 | 0.00001 | 0.00001 | 0.00001 | 0.00001 | 0.00012 | 0.00002 | 0.00003 | 0.00002 | 0.00000 | 0.00003 |
| Clostridium kluyveri | 0.00004 | 0.00002 | 0.00004 | 0.00004 | 0.00003 | 0.00002 | 0.00003 | 0.00003 | 0.00002 | 0.00002 | 0.00003 | 0.00003 | 0.00002 | 0.00012 | 0.00003 | 0.00005 | 0.00006 | 0.00002 | 0.00006 |
| Clostridium ljungdahlii | 0.00000 | 0.00000 | 0.00000 | 0.00000 | 0.00000 | 0.00000 | 0.00000 | 0.00000 | 0.00000 | 0.00000 | 0.00000 | 0.00000 | 0.00000 | 0.00001 | 0.00000 | 0.00000 | 0.00000 | 0.00000 | 0.00000 |
| Clostridium novyi | 0.00003 | 0.00002 | 0.00002 | 0.00005 | 0.00003 | 0.00001 | 0.00003 | 0.00004 | 0.00001 | 0.00003 | 0.00002 | 0.00003 | 0.00001 | 0.00024 | 0.00003 | 0.00004 | 0.00003 | 0.00001 | 0.00004 |
| Clostridium pasteurianum | 0.00005 | 0.00004 | 0.00004 | 0.00008 | 0.00005 | 0.00003 | 0.00005 | 0.00008 | 0.00008 | 0.00005 | 0.00004 | 0.00008 | 0.00003 | 0.00022 | 0.00006 | 0.00007 | 0.00007 | 0.00002 | 0.00009 |
| Clostridium perfringens | 0.00013 | 0.00006 | 0.00009 | 0.00021 | 0.00011 | 0.00006 | 0.00007 | 0.00026 | 0.00012 | 0.00010 | 0.00018 | 0.00007 | 0.00005 | 0.00045 | 0.00009 | 0.00013 | 0.00012 | 0.00011 | 0.00015 |
| Clostridium saccharobutylicum | 0.00003 | 0.00002 | 0.00002 | 0.00005 | 0.00003 | 0.00003 | 0.00003 | 0.00004 | 0.00002 | 0.00002 | 0.00002 | 0.00001 | 0.00001 | 0.00011 | 0.00003 | 0.00001 | 0.00003 | 0.00000 | 0.00005 |
| Clostridium saccharoperbutylaceton | 0.00005 | 0.00003 | 0.00004 | 0.00008 | 0.00004 | 0.00002 | 0.00004 | 0.00005 | 0.00001 | 0.00003 | 0.00004 | 0.00005 | 0.00001 | 0.00017 | 0.00004 | 0.00006 | 0.00004 | 0.00002 | 0.00006 |
| Clostridium scatologenes | 0.00001 | 0.00001 | 0.00000 | 0.00001 | 0.00001 | 0.00000 | 0.00001 | 0.00001 | 0.00000 | 0.00001 | 0.00001 | 0.00001 | 0.00000 | 0.00003 | 0.00001 | 0.00001 | 0.00001 | 0.00000 | 0.00002 |
| Clostridium septicum | 0.00001 | 0.00001 | 0.00001 | 0.00003 | 0.00002 | 0.00001 | 0.00001 | 0.00006 | 0.00001 | 0.00001 | 0.00002 | 0.00001 | 0.00001 | 0.00014 | 0.00001 | 0.00004 | 0.00002 | 0.00000 | 0.00002 |
| Clostridium sp. AWRP | 0.00002 | 0.00001 | 0.00003 | 0.00003 | 0.00001 | 0.00001 | 0.00002 | 0.00002 | 0.00000 | 0.00002 | 0.00001 | 0.00002 | 0.00001 | 0.00004 | 0.00002 | 0.00004 | 0.00003 | 0.00001 | 0.00003 |
| Clostridium sp. BNL1100 | 0.00005 | 0.00003 | 0.00002 | 0.00007 | 0.00004 | 0.00002 | 0.00005 | 0.00005 | 0.00002 | 0.00003 | 0.00003 | 0.00004 | 0.00002 | 0.00011 | 0.00004 | 0.00001 | 0.00007 | 0.00001 | 0.00007 |
| Clostridium sp. CT4 | 0.00002 | 0.00002 | 0.00003 | 0.00004 | 0.00002 | 0.00001 | 0.00004 | 0.00003 | 0.00001 | 0.00003 | 0.00002 | 0.00004 | 0.00001 | 0.00011 | 0.00002 | 0.00002 | 0.00004 | 0.00001 | 0.00004 |
| Clostridium sp. DL-VIII | 0.00006 | 0.00002 | 0.00003 | 0.00006 | 0.00003 | 0.00003 | 0.00006 | 0.00004 | 0.00001 | 0.00004 | 0.00003 | 0.00005 | 0.00001 | 0.00014 | 0.00003 | 0.00002 | 0.00007 | 0.00002 | 0.00006 |
| Clostridium sp. JN-1 | 0.00002 | 0.00001 | 0.00002 | 0.00003 | 0.00002 | 0.00000 | 0.00002 | 0.00001 | 0.00001 | 0.00002 | 0.00002 | 0.00001 | 0.00001 | 0.00010 | 0.00001 | 0.00002 | 0.00004 | 0.00001 | 0.00004 |
| Clostridium sp. JN-9 | 0.00004 | 0.00001 | 0.00002 | 0.00005 | 0.00003 | 0.00001 | 0.00005 | 0.00004 | 0.00001 | 0.00003 | 0.00002 | 0.00003 | 0.00002 | 0.00010 | 0.00002 | 0.00001 | 0.00005 | 0.00001 | 0.00006 |
| Clostridium sp. JN500901 | 0.00001 | 0.00001 | 0.00001 | 0.00002 | 0.00002 | 0.00000 | 0.00001 | 0.00003 | 0.00001 | 0.00001 | 0.00001 | 0.00001 | 0.00000 | 0.00011 | 0.00002 | 0.00003 | 0.00002 | 0.00000 | 0.00003 |
| Clostridium sp. MF28 | 0.00000 | 0.00000 | 0.00000 | 0.00000 | 0.00000 | 0.00000 | 0.00000 | 0.00000 | 0.00000 | 0.00000 | 0.00000 | 0.00000 | 0.00000 | 0.00000 | 0.00000 | 0.00000 | 0.00000 | 0.00000 | 0.00000 |
| Clostridium sp. SYSU GA15002 | 0.00004 | 0.00002 | 0.00002 | 0.00007 | 0.00003 | 0.00002 | 0.00003 | 0.00004 | 0.00001 | 0.00004 | 0.00003 | 0.00003 | 0.00001 | 0.00012 | 0.00003 | 0.00001 | 0.00008 | 0.00001 | 0.00008 |
| Clostridium taeniosporum | 0.00001 | 0.00001 | 0.00001 | 0.00003 | 0.00002 | 0.00001 | 0.00003 | 0.00003 | 0.00001 | 0.00002 | 0.00001 | 0.00003 | 0.00000 | 0.00009 | 0.00001 | 0.00001 | 0.00003 | 0.00001 | 0.00003 |
| Clostridium tetani | 0.00003 | 0.00002 | 0.00002 | 0.00003 | 0.00001 | 0.00001 | 0.00003 | 0.00003 | 0.00001 | 0.00002 | 0.00002 | 0.00003 | 0.00000 | 0.00014 | 0.00002 | 0.00006 | 0.00002 | 0.00000 | 0.00003 |
| Clostridium tyrobutyricum | 0.00001 | 0.00002 | 0.00002 | 0.00002 | 0.00002 | 0.00000 | 0.00002 | 0.00002 | 0.00001 | 0.00001 | 0.00001 | 0.00001 | 0.00001 | 0.00008 | 0.00002 | 0.00001 | 0.00003 | 0.00001 | 0.00002 |
| Clostridium sporogenes | 0.00000 | 0.00000 | 0.00000 | 0.00000 | 0.00000 | 0.00000 | 0.00000 | 0.00000 | 0.00000 | 0.00000 | 0.00000 | 0.00000 | 0.00000 | 0.00000 | 0.00000 | 0.00000 | 0.00000 | 0.00000 | 0.00000 |
| Clostridium;NA | 0.00023 | 0.00013 | 0.00017 | 0.00028 | 0.00022 | 0.00010 | 0.00021 | 0.00032 | 0.00008 | 0.00018 | 0.00015 | 0.00021 | 0.00006 | 0.00114 | 0.00018 | 0.00023 | 0.00029 | 0.00008 | 0.00032 |

| | Subject | | | | | | | | | | | | | | | | | | |
|---|---|---|---|---|---|---|---|---|---|---|---|---|---|---|---|---|---|---|---|
| Blautia | s1 | s2 | s3 | s4 | s5 | s6 | s7 | s8 | s9 | s10 | s11 | s12 | s13 | s14 | s15 | s16 | s17 | s18 | s19 |
| **Blautia sp. SC05B48** | **0.0405** | **0.0138** | **0.0181** | **0.0870** | **0.0374** | **0.0144** | **0.0164** | **0.0175** | **0.0022** | **0.0100** | **0.0104** | **0.0111** | **0.0036** | **0.0273** | **0.0092** | **0.0026** | **0.0423** | **0.0112** | **0.0680** |
| Blautia hansenii | 0.0017 | 0.0011 | 0.0015 | 0.0028 | 0.0022 | 0.0006 | 0.0021 | 0.0031 | 0.0011 | 0.0008 | 0.0019 | 0.0017 | 0.0007 | 0.0025 | 0.0012 | 0.0005 | 0.0024 | 0.0101 | 0.0022 |
| Blautia producta | 0.0026 | 0.0009 | 0.0018 | 0.0022 | 0.0027 | 0.0009 | 0.0043 | 0.0033 | 0.0062 | 0.0018 | 0.0039 | 0.0027 | 0.0021 | 0.0026 | 0.0026 | 0.0038 | 0.0023 | 0.0032 | 0.0038 |
| Blautia sp. N6H1-15 | 0.0027 | 0.0010 | 0.0011 | 0.0040 | 0.0017 | 0.0007 | 0.0026 | 0.0039 | 0.0011 | 0.0007 | 0.0022 | 0.0020 | 0.0008 | 0.0022 | 0.0016 | 0.0007 | 0.0020 | 0.0013 | 0.0020 |
| Blautia sp. YL58 | 0.0000 | 0.0000 | 0.0000 | 0.0000 | 0.0000 | 0.0000 | 0.0000 | 0.0000 | 0.0000 | 0.0000 | 0.0000 | 0.0000 | 0.0000 | 0.0000 | 0.0000 | 0.0000 | 0.0000 | 0.0000 | 0.0000 |
| Blautia;NA | 0.0015 | 0.0008 | 0.0010 | 0.0022 | 0.0014 | 0.0006 | 0.0026 | 0.0020 | 0.0014 | 0.0009 | 0.0018 | 0.0014 | 0.0007 | 0.0022 | 0.0013 | 0.0010 | 0.0020 | 0.0013 | 0.0020 |

| | Subject | | | | | | | | | | | | | | | | | | |
|---|---|---|---|---|---|---|---|---|---|---|---|---|---|---|---|---|---|---|---|
| Eggerthella | s1 | s2 | s3 | s4 | s5 | s6 | s7 | s8 | s9 | s10 | s11 | s12 | s13 | s14 | s15 | s16 | s17 | s18 | s19 |
| **Eggerthella lenta** | **0.0007** | **0.0009** | **0.0011** | **0.0028** | **0.0009** | **0.0046** | **0.0020** | **0.0012** | **0.0129** | **0.0027** | **0.0042** | **0.0011** | **0.0002** | **0.0004** | **0.0044** | **0.0039** | **0.0031** | **0.0007** | **0.0010** |
| Eggerthella sp. YY7918 | 0.0000 | 0.0000 | 0.0000 | 0.0001 | 0.0000 | 0.0000 | 0.0000 | 0.0000 | 0.0000 | 0.0001 | 0.0000 | 0.0000 | 0.0001 | 0.0002 | 0.0001 | 0.0000 | 0.0002 | 0.0000 | 0.0001 |
| Eggerthella;NA | 0.0000 | 0.0000 | 0.0000 | 0.0000 | 0.0000 | 0.0000 | 0.0000 | 0.0000 | 0.0000 | 0.0000 | 0.0000 | 0.0000 | 0.0000 | 0.0000 | 0.0000 | 0.0000 | 0.0000 | 0.0000 | 0.0000 |

| | Subject | | | | | | | | | | | | | | | | | | |
|---|---|---|---|---|---|---|---|---|---|---|---|---|---|---|---|---|---|---|---|
| Barnesiella | s1 | s2 | s3 | s4 | s5 | s6 | s7 | s8 | s9 | s10 | s11 | s12 | s13 | s14 | s15 | s16 | s17 | s18 | s19 |
| **Barnesiella viscericola** | **0.00173** | **0.00061** | **0.00181** | **0.00115** | **0.00062** | **0.00132** | **0.00118** | **0.00049** | **0.00023** | **0.00097** | **0.00069** | **0.00096** | **0.00201** | **0.00198** | **0.00123** | **0.00094** | **0.00066** | **0.00052** | **0.00097** |

| | Subject | | | | | | | | | | | | | | | | | | |
|---|---|---|---|---|---|---|---|---|---|---|---|---|---|---|---|---|---|---|---|
| Butyricimonas | s1 | s2 | s3 | s4 | s5 | s6 | s7 | s8 | s9 | s10 | s11 | s12 | s13 | s14 | s15 | s16 | s17 | s18 | s19 |
| **Butyricimonas faecalis** | **0.0016** | **0.0041** | **0.0012** | **0.0029** | **0.0018** | **0.0056** | **0.0007** | **0.0020** | **0.0006** | **0.0034** | **0.0049** | **0.0043** | **0.0062** | **0.0076** | **0.0013** | **0.0041** | **0.0027** | **0.0012** | **0.0012** |

| | Subject | | | | | | | | | | | | | | | | | | |
|---|---|---|---|---|---|---|---|---|---|---|---|---|---|---|---|---|---|---|---|
| Collinsella | s1 | s2 | s3 | s4 | s5 | s6 | s7 | s8 | s9 | s10 | s11 | s12 | s13 | s14 | s15 | s16 | s17 | s18 | s19 |
| **Collinsella aerofaciens** | **0.04814** | **0.02807** | **0.01038** | **0.02709** | **0.01433** | **0.03724** | **0.05531** | **0.0018** | **3.2E-05** | **0.00023** | **8E-05** | **0.00045** | **0.00011** | **0.04884** | **0.00088** | **0.00029** | **0.02341** | **0.00023** | **0.00144** |

| | Subject | | | | | | | | | | | | | | | | | | |
|---|---|---|---|---|---|---|---|---|---|---|---|---|---|---|---|---|---|---|---|
| Faecalibacterium | s1 | s2 | s3 | s4 | s5 | s6 | s7 | s8 | s9 | s10 | s11 | s12 | s13 | s14 | s15 | s16 | s17 | s18 | s19 |
| **Faecalibacterium prausnitzii** | **0.0751** | **0.0288** | **0.0832** | **0.0585** | **0.0641** | **0.0253** | **0.0773** | **0.0960** | **0.0110** | **0.0279** | **0.0026** | **0.0870** | **0.0256** | **0.1414** | **0.0799** | **0.0057** | **0.0189** | **0.0451** | **0.0986** |

| | Subject | | | | | | | | | | | | | | | | | | |
|---|---|---|---|---|---|---|---|---|---|---|---|---|---|---|---|---|---|---|---|
| Odoriobacter | s1 | s2 | s3 | s4 | s5 | s6 | s7 | s8 | s9 | s10 | s11 | s12 | s13 | s14 | s15 | s16 | s17 | s18 | s19 |
| **Odoribacter splanchnicus** | **0.0054** | **0.0054** | **0.0058** | **0.0043** | **0.0055** | **0.0070** | **0.0090** | **0.0048** | **0.0002** | **0.0046** | **0.0027** | **0.0113** | **0.0124** | **0.0082** | **0.0048** | **0.0036** | **0.0061** | **0.0020** | **0.0049** |

Dark green indicates greater abundance. If multiple bacterial strains were detected by shotgun metagenomic analysis within a single genus, the strain with the highest abundance among all participants was selected for single regression analysis.

**Supplementary Table 8. Nonsignificant Results of Regression Analysis for Intestinal Microbiota and Physical and Psychological Indices (Study 2)**

| | *β* | *std.Error* | *t - value* | *p* | $R^2$ | *F-value* |
|---|---|---|---|---|---|---|
| Barnesiella viscericola × CVI | 0.04 | 0.27 | 0.14 | 0.89 | -0.03 | $F_{1, 16}$ = 0.53 |
| Blautia SC05B48 × PSI-parent | -1.71 | 10.50 | -0.16 | 0.87 | -0.03 | $F_{1, 17}$ = 0.48 |
| Butyricimonas faecalis × CVI | 0.28 | 0.20 | 1.42 | 0.18 | 0.05 | $F_{1, 15}$ = 1.86 |
| Butyricimonas faecalis × oxytocin | 9.10 | 74.39 | 0.12 | 0.90 | 0.06 | $F_{1, 15}$ = 2.03 |
| Collinsella aerofaciens × CSI | 0.12 | 0.17 | 0.73 | 0.48 | 0.07 | $F_{1, 17}$ = 2.43 |
| Eggerthella lenta × CSI | -0.11 | 0.41 | -0.26 | 0.80 | 0.02 | $F_{1, 17}$ = 1.41 |
| Faecalibacterium prausnitzii × maximum gait speed | 0.04 | 0.11 | 0.37 | 0.72 | -0.05 | $F_{1, 17}$ = 0.13 |
| Odoribacter splanchnicus × maximum gait speed | -0.05 | 0.14 | -0.35 | 0.73 | -0.05 | $F_{1, 17}$ = 0.12 |
| Odoribacter splanchnicus × J-RS | 6.77 | 4.55 | 1.49 | 0.16 | 0.05 | $F_{1, 17}$ = 1.98 |

Significant relationships are shown in Figure 6 in the main text.

*β* = standardized partial regression coefficient.
